# EF-P and its paralog EfpL (YeiP) differentially control translation of proline-containing sequences

**DOI:** 10.1101/2024.04.15.589488

**Authors:** Alina Sieber, Marina Parr, Julian von Ehr, Karthikeyan Dhamotharan, Pavel Kielkowski, Tess Brewer, Anna Schäpers, Ralph Krafczyk, Fei Qi, Andreas Schlundt, Dmitrij Frishman, Jürgen Lassak

## Abstract

Polyproline sequences (XPPX) stall ribosomes, thus being deleterious for all living organisms. In bacteria, translation elongation factor P (EF-P) plays a crucial role in overcoming such arrests. 12% of eubacteria possess an EF-P paralog – YeiP (EfpL) of unknown function. Here, we functionally and structurally characterize EfpL from Escherichia coli and demonstrate its yet unrecognized role in the translational stress response. Through ribosome profiling, we analyzed the EfpL arrest motif spectrum and discovered additional stalls beyond the canonical XPPX motifs at single-proline sequences (XPX), that both EF-P and EfpL can resolve. Notably, the two factors can also induce pauses. We further report that, contrary to the housekeeping EF-P, EfpL can sense the metabolic state of the cell, via lysine acylation. Together, our work uncovers a new player in ribosome rescue at proline-containing sequences, and provides evidence that co-occurrence of EF-P and EfpL is an evolutionary driver for higher bacterial growth rates.

## Introduction

Decoding genetic information at the ribosome is a fundamental trait shared among all living organisms. However, translation of two or more consecutive prolines leads to ribosome arrest^1–6^. To allow translation to continue, nearly every living cell is equipped with a specialized elongation factor called e/aIF-5A in eukaryotes and archaea, or EF-P in bacteria^7,8^. Upon binding close to the ribosomal tRNA exiting site (E-site), EF-P stimulates peptide bond formation by stabilizing and orienting the peptidyl-tRNA^Pro^ ^9,10^. EF-P has a three-domain structure that spans both ribosomal subunits^10,11^ and consists of an N-terminal KOW domain and two OB domains^12^, together mimicking tRNA in size and shape^13^. Although this structure is conserved among all EF-P homologs^14^, bacteria have evolved highly diverse strategies to facilitate proper interactions between EF-P and the CCA end of the P-site tRNA^Pro^. For instance, in *Escherichia coli* and 25% of all bacteria a conserved lysine K34 at the tip of the loop bracketed by two beta strands β3 and β4 (β3Ωβ4) is post-translationally activated by β-D-lysylation using the enzyme EpmA^15–19^. Firmicutes such as *Bacillus subtilis* elongate lysine K32 of their EF-P by 5-aminopentanolylation^20^, while in about 10% of bacteria, including pseudomonads, an arginine is present in the equivalent position, which is α-rhamnosylated by the glycosyltransferase EarP^14,21,22^. Among the remaining EF-P subtypes the paralogous YeiP (from now on termed EfpL for ‘EF-P like’) sticks out, as it forms a highly distinct phylogenetic branch (**Fig. 1A; Extended Data Fig. 1**) ^14,23^ suggesting that its role in translation diverges from those of canonical EF-Ps. However, to date the molecular function of EfpL remains enigmatic. In the frame of this study, we solved the structure of *E. coli* EfpL (EfpL) and uncovered its role in translation of XP(P)X-containing proteins.

**Fig. 1:**
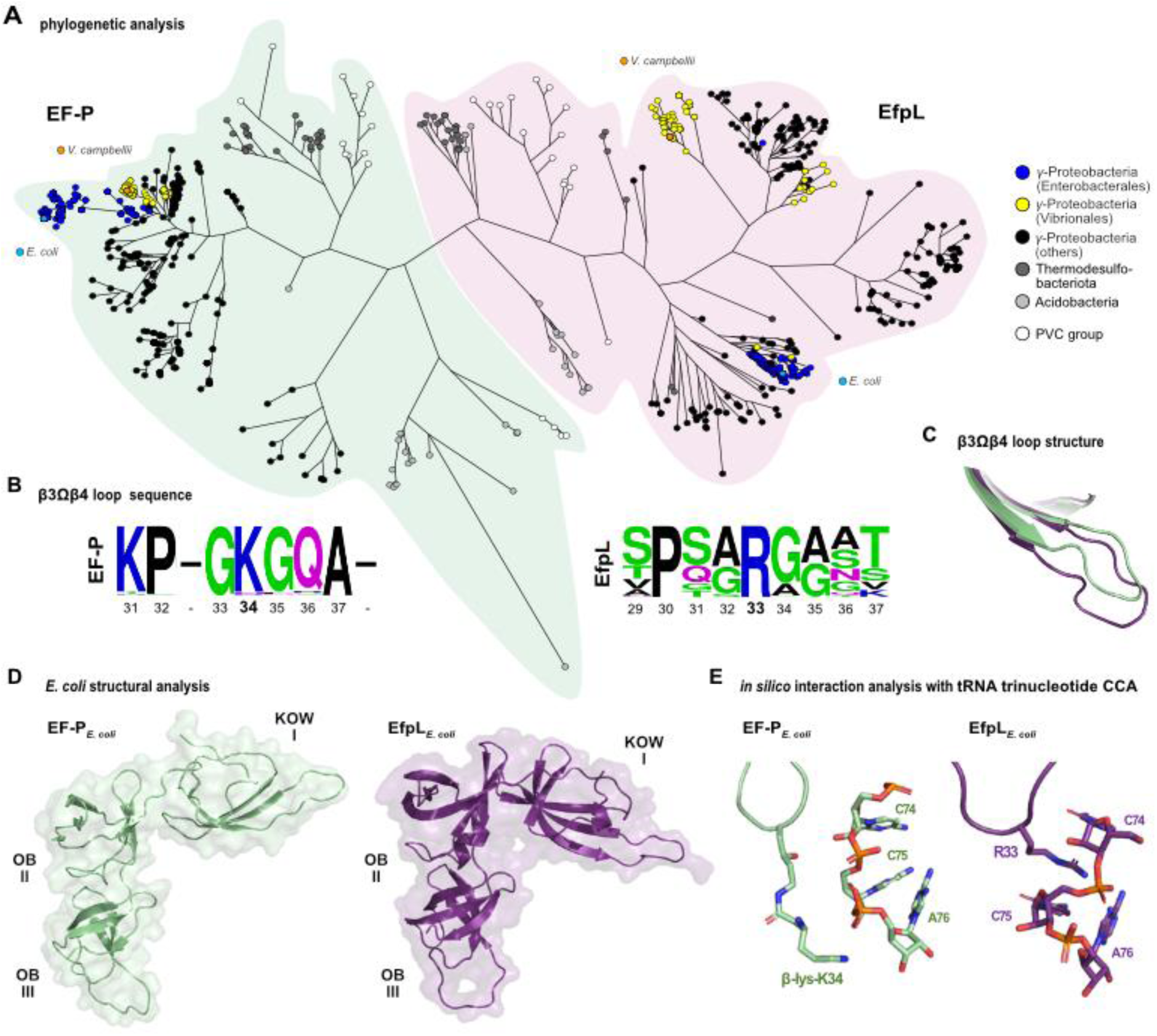
Structural and phylogenetic analysis of the EfpL subgroup. **(A)** Phylogenetic tree of EfpL (purple) and co-occurring EF-Ps (green). Colors of tip ends depict bacterial clades. **(B)** Sequence logos^25^ of β3Ωβ4 loop of EfpL and co-occurring EF-Ps. **(C)** Comparison of the KOW β3Ωβ4 loop in *E. coli* EF-P (taken from PDB: 6ENU; green) and EfpL (PDB: 8S8U, this study; purple). **(D)** Comparison of structures of *E. coli* EF-P (taken from PDB: 6ENU) and EfpL (PDB: 8S8U, this study) with overall fold views and three domains. **(E)** Excerpt of β3Ωβ4 loops from EF-P and EfpL in complex with the tRNA trinucleotide CCA. The central tip residue of EF-P is β-lysylated.

## Results

### Structural and phylogenetic analysis of EfpL revealed unique features in the β3Ωβ4 loop

We began our study by recapitulating a phylogenetic tree of EF-P in order to extract the molecular characteristics of the EfpL subgroup. A collection of 4736 complete bacterial genomes was obtained from the RefSeq database^24^. From these organisms, we extracted 5448 EF-P homologs and identified the branch that includes the “elongation factor P-like protein” YeiP of *E. coli.* This subfamily comprises 528 sequences (**Extended Data Fig. 1; Supplementary Fig. S1; Supplementary Tab. S1**) and is characterized by a number of unique features (**Fig. 1**). First, we observed that EfpL is predominantly found in Proteobacteria of the γ-subdivision but also in Thermodesulfobacteria, Acidobacteria and the Planctomycetes/Verrucomicrobia/Chlamydiae-group (PVC-group) (**Fig 1A**). The protein strictly cooccurs with a canonical, mostly lysine-type, EF-P (**Supplementary Fig. S1A, B**), suggesting a similar but more specialized role in translation than EF-P. Second, we noted that the EfpL branch is most closely affiliated but still separated from the EF-P subgroup which is activated by EarP (**Extended Data Fig. 1**). This evolutionary connection extends beyond overall sequence similarity to the functionally significant β3Ωβ4 loop and the arginine (R33 in EfpL) at its tip (**Fig 1B; Supplementary Fig. S1C**) ^14^. However, in contrast to EarP-type EF-Ps, R33 in EfpL remains unmodified, as confirmed by mass spectrometry (MS) (**Supplementary Fig. S2**). Additionally, we discovered a strictly conserved proline three amino acids upstream of EfpL_R33 – an amino acid typically absent from that position in EarP-type EF-Ps^14^. Third, EfpLs predominantly co-occur with the EF-P subfamily activated by EpmA whereas the presence of an EarP-type *efp* in the genome typically excludes the existence of the paralogous EfpL (**Supplementary Fig. S1C**) ^23^. Lastly, distinguishing itself from all other EF-Ps, EfpL appears to possess a β3Ωβ4 extension (**Fig. 1B, C; Supplementary Fig. S1D**). However, the exact length of this extension remains ambiguous in the *in silico* models.

Accordingly, we solved the crystal structure of *E. coli* EfpL (**PDB: 8S8U; Supplementary Tab. S2A**) and compared it with other available protein structures of EF-P^10,26^. This confirmed the highly conserved fold of EF-P typed proteins in prokaryotes, both expressed by a structural overlay and respective root-mean-square deviation (r.m.s.d.) values (**Extended Data Fig. 2**). Interestingly, the EfpL structure reveals a significantly tilted KOW domain relative to the C-terminal di-domain compared to EF-P structures (**Fig. 1D**), certainly enabled by the flexible hinge region between the independent moieties. However, a separate alignment of KOW and OB di-domains between *E. coli* EfpL and for example the EF-P structure resolved within the *E. coli* ribosome from Huter *et al*. ^10^, reveals low r.m.s.d. values (**Extended Data Fig. 2**). This suggests the relative domain arrangement is merely a consequence of the unique crystal packing. Altogether, the EfpL high-resolution structure reveals the anticipated fold and features needed for its expected functional role interacting with the ribosome, analogously to EF-P.

We then took a closer look at the KOW domain β3Ωβ4 loop relevant for interacting with the tRNA. The structural alignment ultimately revealed a β3Ωβ4 loop elongation by two amino acids for EfpL, different from the canonical seven amino acids in EF-P (**Fig. 1B, C**). In this way, EfpL_R33 remains apical similar to canonical EF-Ps. We reasoned that such a loop extension would enable unprecedented contacts with the CCA end of the P-site tRNA without further post-translational modification, which we set out to investigate in detail. Given the overall structural similarity with EF-P we overlaid the EfpL KOW-domain with the cryo-EM structure of EF-P bound to the ribosome^10^ to analyze the position and potential contacts of EfpL_R33 with the mRNA trinucleotide. In EF-P, the modified K34 aligns with the trinucleotide backbone without obvious RNA-specific interactions, while the prolonged sidechain allows for a maximum contact site with the RNA. To allow for local adjustments in an otherwise sterically constrained frame of the ribosome, we carried out molecular docking of EfpL and the trinucleotide with a local energy minimization using HADDOCK^27^ (**Fig. 1E; Extended Data Fig. 3; Supplementary Tab. S2B**). As shown in the lowest-energy model, we find that the unmodified arginine in the EfpL β3Ωβ4 loop can reestablish the interaction. Furthermore, EfpL can mediate specific interactions with the RNA as – unlike EF-P_K34 – R33 stacks between the two C-bases and makes polar interactions with the phosphate-sugar backbone. We thus conclude that the prolonged β3Ωβ4 loop and its central tip R33 are capable of compensating for the lack of a modified lysine.

### *E. coli* EF-P and EfpL have overlapping functions

Based on the structural similarities (**Fig. 1D**), we assumed that EF-P and EfpL have a similar molecular function. However, there has been no experimental evidence supporting this hypothesis so far. Accordingly, we analyzed growth of *E. coli* wild type and mutants lacking *efp* (Δ*efp*), *efpL* (Δ*efpL*), or both genes (Δ*efp* Δ*efpL*) (**Fig. 2A**). Compared to the strong mutant phenotype in Δ*efp* (t_d_ ∼31 min), we observed a slight but still significant increase in doubling time from ∼20 min in the wild type to ∼24 min in Δ*efpL*. Deletion of both proteins in Δ*efp* Δ*efpL* impairs growth beyond the loss of each single gene (t_d_ ∼40min). This implies a cooperative role in the translation of polyproline proteins, which is almost masked by EF-P in Δ*efpL* cells. The overproduction of either EF-P or EfpL, but not the substitution of the functional important R33 at the β3Ωβ4 loop tip in the EfpL_R33K variant, completely or partially eliminates the growth defect. However, this effect vanishes when the functional important R33 at the β3Ωβ4 loop tip is substituted in the EfpL_R33K variant, demonstrating the significance of R33 for the molecular function of EfpL. It is also noteworthy that overproduction of EfpL in Δ*efp* Δ*efpL* reduced doubling time below that of Δ*efp* (∼27 min). We hypothesize that ectopic expression compensates for the comparative low copy number of EfpL per cell (EfpL: ∼4500 vs. EF-P: ∼40,000 in complex medium^28^) (**Supplementary Fig. S3**). A similar phenotypic pattern emerges when examining the same strains in terms of the CadABC-dependent pH stress response (**Supplementary Fig. S4**) ^29^, whose regulator CadC has a polyproline motif^3^. Next, we investigated EfpL’s capability to relieve ribosome arrest on diprolines. In an *in vivo* approach, we used our recently described reporter assay (**Fig. 2B**) ^30^. This assay allows positive correlation of translational pausing strength with bioluminescence. Deletion of either *efp* or *efpL* leads to an increased light emission, and for Δ*efp* Δ*efpL*, we observed a cumulative effect (**Fig. 2C**). Again, the phenotype of Δ*efp* Δ*efpL* was trans-complemented by wild-type copies of the respective genes. A parallel quantitative *in vitro* assay employing NanoLuc® variants with and without polyproline insertion (**Fig. 2D**) confirmed the results of the previous *in vivo* experiments with EfpL and its substitution variant EfpL_R33K (**Fig. 2E**). Interestingly, unlike in the *in vivo* analyses with Δ*efpL* and Δ*efp* strains, there are no significant differences in the rescue efficiency between EF-P and EfpL at the tested diproline motif PPN.

**Fig. 2:**
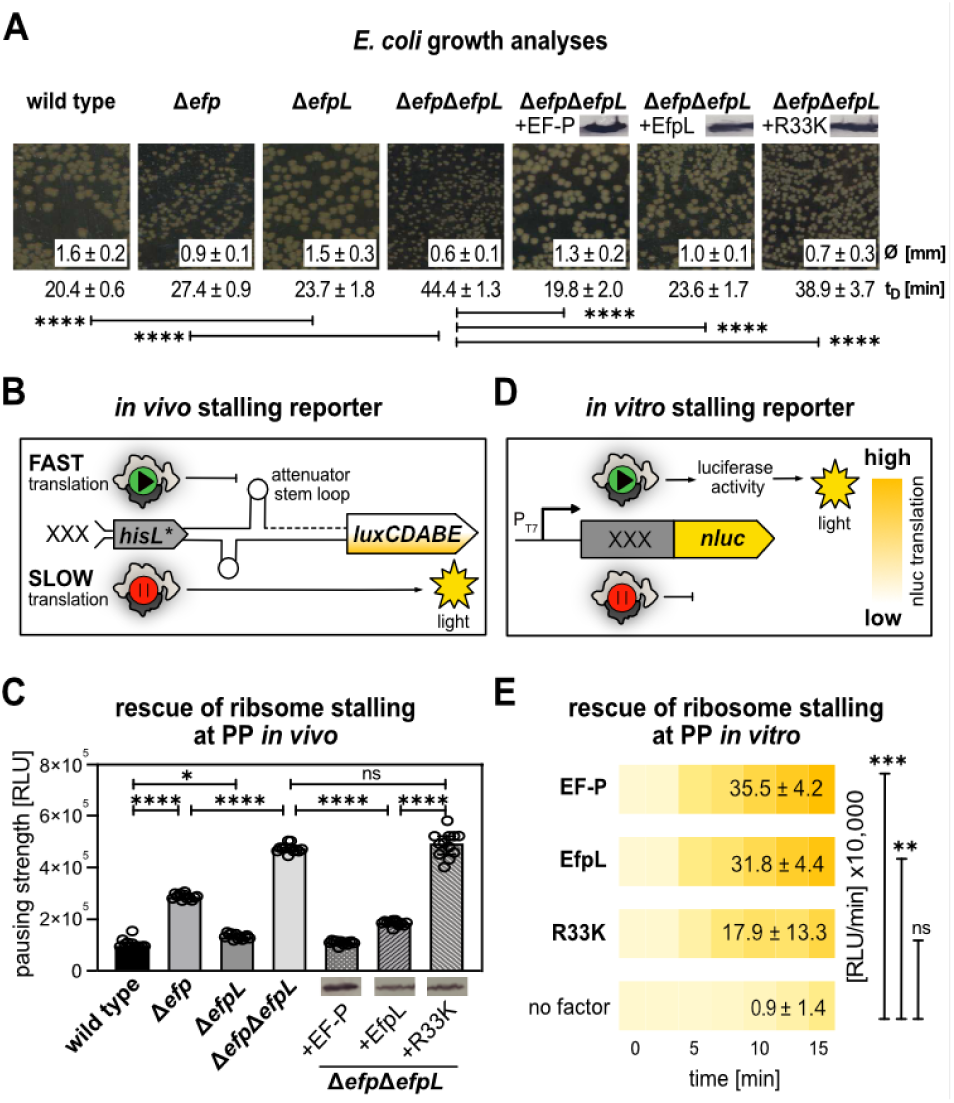
The role of EfpL in bacterial physiology. **(A)** Growth analysis of *E. coli* BW23113 and isogenic mutant strains lacking *efp (*Δ*efp)*, *efpL (*Δ*efpL)*, or both genes (Δ*efp*Δ*efpL)*. For complementation gene copies of *efp* (+EF- P), *efpL* (+EfpL) or *efpL_R33K* (+EfpL_R33K) were provided in trans. Protein production was confirmed by immunoblotting utilizing the C-terminally attached His_6_-tag and Anti-His_6_ antibodies (α-His). Colony size was quantified by averaging the diameters (∅ ± standard deviation) of 30 colonies on LB agar plates after 18h of cultivation at 37°C. Doubling times (t_D_ ± standard deviation) were calculated from exponentially grown cells in LB (n = 12). Statistically significant differences according to ordinary one-way ANOVA test (*P value <0.0332, **P value <0.0021, ***P value <0.0002, ****P value <0.0001, ns not significant). **(B)** Scheme of the *in vivo* stalling reporter system ^30^. The system operates on the histidine biosynthesis operon of *E. coli*. In its natural form, the histidine biosynthesis gene cluster is controlled by the His-leader peptide (HisL), which comprises seven consecutive histidines. In our setup, the original histidine residues (His1 through His4) were replaced by artificial sequence motifs (XXX). Non-stalling sequences promote the formation of an attenuator stem loop (upper part) that impedes transcription of the downstream genes, thus ultimately preventing light emission. Conversely, in the presence of an arrest motif, ribosomes pause and hence an alternative stem loop is formed that does not attenuate transcription of the *luxCDABE* genes of *Photorhabdus luminescens*. **(C)** *In vivo* comparison of pausing at PPN in *E. coli* (for strain labeling and immunoblotting details see (A)). Pausing strength is given in relative light units (RLU) (n = 12, Error bars indicate standard deviation). Statistics were determined and labeled as in (A). **(D)** Scheme of the *in vitro* cell-free stalling reporter assay. The system is based on *nanoluc* luciferease (nluc®) which is preceded by an artificial sequence motif (XXX). DNA is transcribed from a T7 promoter (P_T7_) using purified T7 polymerase (NEB). Pausing strength is proportional to light emission. **(E)** *In vitro* transcription and translation of the nLuc® variants nLuc_PPN. The absence (no factor) or presence of the respective translation elongation factors of *E. coli* (EF-P, EfpL, EfpL_R33K) is shown. Translational output was determined by measuring bioluminescence in a time course of 15 minutes and endpoints are given in relative light units (RLU/min) (n ≥ 3). Statistics were determined and labeled as in (A).

### *E. coli* EF-P and EfpL alleviate ribosome stalling at distinct XP(P)X motifs with differences in rescue efficiency

To elucidate the EfpL arrest motif spectrum, a ribosome profiling analysis (RiboSeq) was conducted. Here an *E. coli* wild type was compared with Δ*efp* and Δ*efpL* strains. Importantly, we also included Δ*efp* cells in which EfpL was overproduced. As indicated by our previous analyses (**Fig. 2**) this compensates for the low natural copy number of the factor and might uncover motifs that are otherwise masked by the presence of EF-P. We used PausePred^31^ to predict pauses in protein translation in the respective strains. Subsequently, we calculated the frequencies of amino acid triplets residues occurring at the sites of predicted pauses (**Fig. 3A; Supplementary Tab. S3A**). In line with the molecular function of EF-P, diproline motifs were heavily enriched at pause sites in Δ*efp*^3,5,6^. As already suspected by the mild mutant phenotype of the *efpL* deletion (**Fig. 2**) we did not see a significant difference between Δ*efpL* and wild type. However, in stark contrast, overproduction of EfpL alleviated ribosome stalling at many but not all arrest motifs identified in Δ*efp*. This further corroborates the idea that EfpL has evolved to assist EF-P in translational rescue. Most striking in our analysis was the observation that among the top 29 stalling motifs we found not only XPPX, but also many XPX motifs and even one motif completely lacking a proline. The RiboSeq findings were confirmed with our *in vivo* luminescence reporter (**Fig. 3B; Extended Data Fig. 4**) by testing 12 different arrest motifs as well as *in vitro* by quantifying production of two NanoLuc® Luciferase (nLuc) variants comprising IPW and PAP (**Fig. 3C; Extended Data Fig. 5**). Together, our data demonstrates that while a P-site proline is almost always a prerequisite for ribosome rescue by EF-P/EfpL, in rare cases motifs lacking proline can also be targeted.

**Fig 3:**
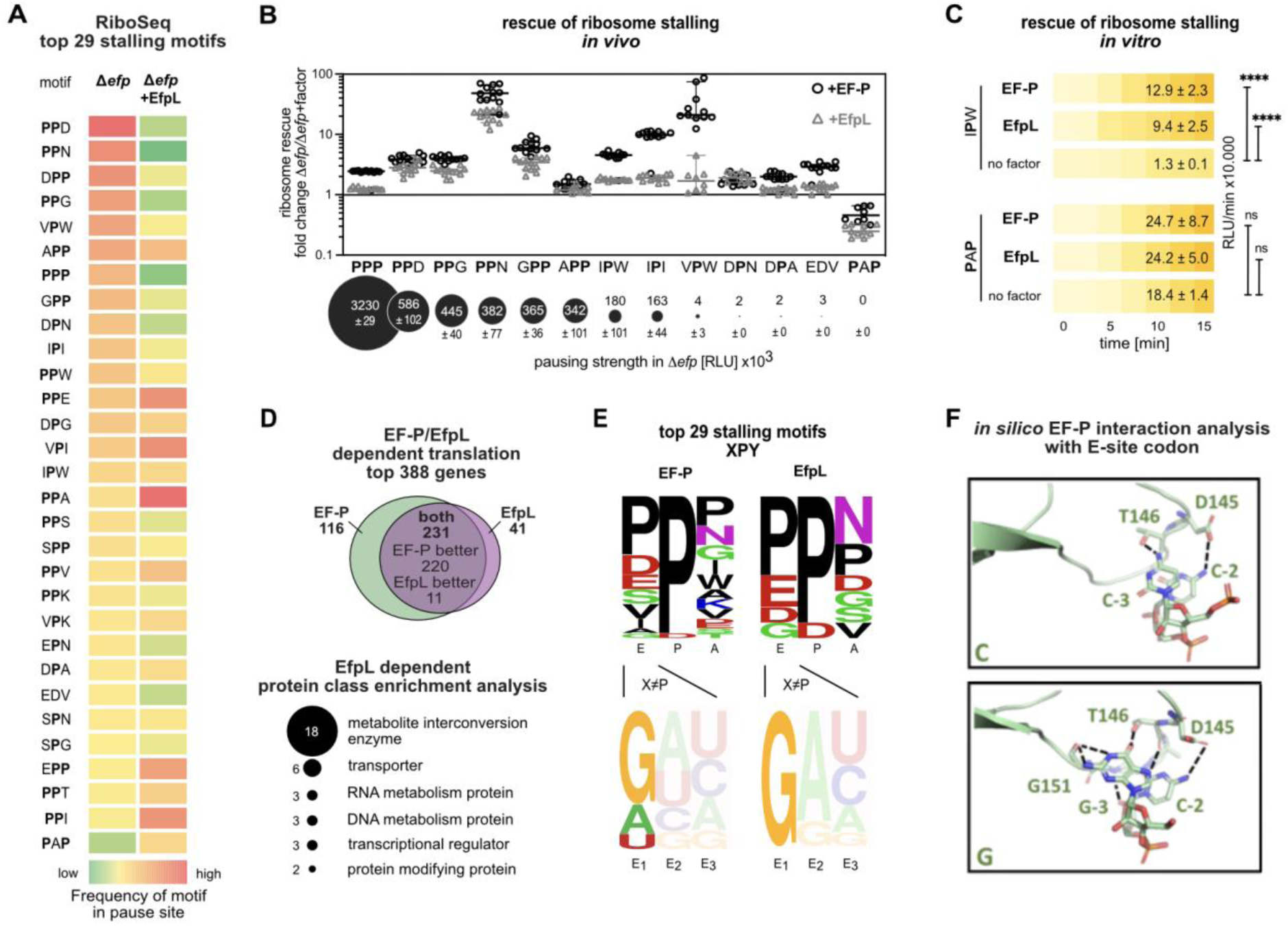
The target spectrum of EF-P and EfpL. **(A)** First column: The top 29 motifs whose translation is dependent on EF-P and the control motif PAP in the ribosome profiling analysis comparing *E. coli* BW25113 wild type with the *efp* deletion mutant (Δ*efp*). Second column: Comparison of ribosome profiling data of Δ*efp* cells and Δ*efp* cells overexpressing *efpL* (Δ*efp* +EfpL) at these motifs. The color code of the heat map corresponds to the the frequency of the motif to occur in pause site predicted with PausePred^31^ (From green to red = from low to high). **(B)** *In vivo* comparison of rescue efficiency of a set of stalling motifs and the control motif PAP. Given is the quotient of relative light units measured in Δ*efp* and corresponding trans complementations by EF-P (+EF-P, open circles) and EfpL (+EfpL, open triangles). Motifs are sorted according to pausing strength determined with our previously introduced stalling reporter. Statistics: n = 12, Error bars indicate standard deviation. Significant differences were determined according to a 2-way ANOVA test (*P value <0.0332, **P value <0.0021, ***P value <0.0002, ****P value <0.0001, ns not significant). **(C)** *In vitro* transcription and translation of the *nLuc®* variants nLuc_3xRIPW (IPW) and nLuc_3xRPAP (PAP). The absence (no factor) or presence of the respective translation elongation factors of *E. coli* (EF-P, EfpL) is shown. Translational output was determined by measuring bioluminescence in a time course of 15 minutes and is given in relative light units measured at the end of the reaction (RLU/min ± standard deviation) (n ≥ 3). Statistically significant differences according to ordinary one-way ANOVA test. **(D)** Upper part: Venn diagram of the top 388 genes whose translation depends on EF-P and EfpL. Dependency was determined by comparing asymmetry scores from genes encompassing the top 29 stalling motifs listed in A). Lower part: Enriched protein classes to which EfpL dependent genes belong^32^. **(E)** Sequence composition combining the top 29 stalling motifs being either EF-P dependent (left) or EfpL dependent (right). Bottom: Sequence logo^25^ of the E-site codon composition of XPY motifs, where X≠P. **(F)** Comparison of EF-P OB domain 3 loop 1 in contact with the E-site tRNA codons _-3_CCG_-1_ or _-3_ GCG_-1_. Only the first two nucleotides are show for clarity.

An arrest spectrum extension beyond diprolines has only been reported for IF-5A thus far^33,34^ although there are indications in the literature that EF-P might assist in synthesis of an XPX containing sequence^35–37^. We confirmed these data for the XPX containing leader peptide MgtL (**Supplementary Fig. S5**) and found that EfpL similarly contributes to alleviate stalling at this sequence. To further explore EfpL’s contribution to gene specific translational rescue, we focused on the top 29 motifs as done before for eIF-5A^33^ and looked at the frequency of ribosome occupancy before and after the pause sequence. The ratio between these values gives an asymmetry score (AS) and provides a good measure for stalling strength^6^. EF-P and EfpL dependency was determined by comparing with the AS from the wild type. We were thus able to recapitulate the data from previous RiboSeq analyses for the Δ*efp* samples (**Supplementary Tab. S3B, C**). Moreover, with this approach we were able to find EfpL targets not only in the Δ*efp* +EfpL sample but also in Δ*efpL*. In line with our phenotypic analysis (**Fig. 1; Supplementary Fig. S1; Supplementary Fig. S4; Supplementary Fig. S5**), most of these proteins are also targeted by EF-P (**Fig. 3D**; **Supplementary Tab. S3**). While in the majority of cases, the rescue efficiency was better with EF-P, we found some proteins where EfpL seems to be superior. We even identified a few candidates that were only dependent on EfpL. The proteins targeted by EfpL are frequently involved in amino acid metabolism and transport (**Fig. 3D; Supplementary Tab. S3D**). This provides a potential explanation for the growth phenotype we observed in Lysogeny broth (LB), where amino acids constitute the major source of nutrients. Taken together our data demonstrate that both EF-P and EfpL jointly regulate the translation of XPPX and XPX proteins and both factors being necessary for maximal growth speed under tested conditions.

### A guanosine in the first position of the E-site codon as recognition element for EF-P and EfpL

The chemical nature of the X residues in XP(P)X in the top 29 stalling motifs (**Fig. 3A, E**) is highly diverse and does not provide a cohesive rationale for the arrest motif spectrum: besides the negatively charged residues aspartate and glutamate, we found especially the hydrophobic amino acids isoleucine and valine as well as small ones, like glycine for X at the XP(P) position. Consequently, we extended our view to the codon level. EF-P and accordingly EfpL can interact with the E-site codon utilizing the first loop in the C-terminal OB-domain (d3 loop I) ^10,11^. We did not see any preference for a specific base in the wobble position. By contrast, we revealed a strong bias for guanosine in the first position of the E-site codon in the sequence logos (**Fig. 3E; Supplementary Fig. S6**) of EF-P- and EfpL- targeted XPP motifs, where X≠P. Notably, we observed no clear trend when we looked at the X in (P)PX in motifs (**Supplementary Fig. S6**). When bound to the ribosome, EF-P establishes contacts with the first and second position of the E-site codon through d3 loop I residues G144–G148, with sidechain-to-base specific contacts involving D145 and T146^10^ (**Fig. 3F**). However, in the available high-resolution structure, ribosomes are arrested at a triproline motif and thus, the E-site codon (CCN) does not contain a guanosine. Referring to our observation we replaced the cytosine in the structure by guanosine *in silico* (**Extended Data Fig. 6**), followed by an additional docking and energy minimization of the loop-RNA interface. The resulting complex supports the appearance of additional contacts possible between guanosine and EF-P compared to cytosine (**Fig. 3F; Extended Data Fig. 6**). This is in particular supported by an extended interface with sequential contacts up to residue G151, thus involving the entire d3 loop I. We therefore conclude that especially guanosine in first position of the E-site codon promotes EF-P and EfpL binding to the ribosome.

### EF-P and EfpL can induce translational pauses

We found the unique recognition elements of an EF-P/EfpL dependent arrest motif to be the P-site tRNA^Pro^ and the E-site codon, in agreement with past studies^9,10,38^. We therefore wondered whether XP – regardless of being part of a stalling motif or not – promotes binding of EF-P and similarly EfpL to the ribosome. If so, such “off-binding” might induce pausing at non-stalling motifs instead of alleviating it. Although weak, we indeed saw that loss of *efp* increases pausing with our PAP non-stalling control (**Fig. 3B**), which comprises two XPX motifs namely RPA and APH. Conversely, *efp* and *efpL* overexpression showed the opposite effect. Thus, our study provides first evidence that the translation factors EF-P and EfpL can induce pausing, presumably by blocking tRNA translocation to the E-site. Our hypothesis was confirmed by showing that one can also induce pausing at a clean APH motif (**Supplementary Fig. S7**). Either such an apparently deleterious effect is accepted, as the positive influence on arrest motifs outweighs the negative one, or translational pauses at XP(P)X might also have positive effects on, for example, buying time for domain folding or membrane insertion^39^. We were further curious whether we see codon specific effects and tested the non-stalling motif RPH, in which the E-site codon starts with C (R is encoded by CGC) (**Supplementary Fig. S7**). Congruent with our previous findings (**Fig. 3E**) EF-P could no longer increase pausing strength and with EfpL the effect was less pronounced. In summary our findings indicate that EF-P (and EfpL) may be able to bind to the ribosome whenever a proline is translated, with binding being further promoted by the E-site codon. This idea is in line with earlier work from Mohapatra *et al.* ^40^. The authors reported that EF-P binds to ribosomes during many or most elongation cycles. Our data may now provide a rationale for this (at the time) unexpectedly high binding frequency, which by far exceeds the number of XPPX arrest motifs. In addition to these weak pauses induced at XPX, we observed in our RiboSeq data that EF-P might also bind non-productively at certain motifs as evidenced by asymmetry scores that are higher in Δ*efp* samples than in the wild type (**Supplementary Tab. S3B, C**). While such events are predominantly weak and only rarely observed in our Δ*efp* RiboSeq data, their frequency and strength increased when we overproduced EfpL in the Δ*efp* +EfpL sample (**Supplementary Fig. S7; Supplementary Tab. S3B, C**). This supports the idea that the structural differences of the two factors differentially align and stabilize the P-site tRNA^Pro^. We thus reasoned that the presence of a constitutive EF-P and a more specialized EfpL, would provide the cell with a lever to intentionally delay or accelerate translation gene specifically. However, this would require regulation. Neither in our data (**Supplementary Fig. S8**) nor in the literature^28^ did we find any hint for transcriptional or translational control of EfpL. Instead, the copy number ratio of EF-P:EfpL was consistent in all tested condition^28^.

### *E. coli* EfpL activity is regulated by multiple acylations in the KOW domain

As an alternative to copy number control, post-translational modifications provide a means to adjust EF-P activity to cellular needs. Since we were able to demonstrate that – unlike many other EF-P subtypes – the EfpL β3Ωβ4 loop tip is unmodified, we extended our view to the entire protein sequence. The idea arose as the activity and subcellular localization of the eukaryotic EF-P ortholog eIF5A is regulated by phosphorylation and acetylation, respectively^41,42^. Strikingly, a literature search revealed that *E. coli* EfpL is acylated at four different lysines (K23, K40, K51 and K57) in the KOW domain (**Fig. 4A**) ^43–46^. Notably, a sequence comparison with EF-P shows that a lysine is found only in the position equivalent to K57, and there is no evidence of modification^43–46^. Possible acylations of EfpL encompass not only acetylation but also malonylation and succinylation (**Fig. 4A**). As a consequence, the positive charge of lysine can either be neutralized or even turned negative. To investigate the impact of acylation on EfpL we generated protein variants in which we introduced *N*_ε_-acetyllysine by amber suppression^47^ at each individual position where acylation was previously reported (EfpL_K23AcK, EfpL_K40AcK, EfpL_K51AcK and EfpL_K57AcK). Testing of purified protein variants in the established *in vitro* assay revealed that K51 acetylation impairs EfpL’s function, significantly (**Fig. 4B; Extended Data Fig. 7**). We argue that charge alterations at these lysines as well as subsequent steric constraints will impair ribosomal interactions. To this end, we modelled the EfpL KOW domain to the ribosome by structural alignment with EF-P in order to investigate the effects of acetylation visualized by respective *in silico* modifications (**Extended Data Fig. 8**). In line with rescue experiments, the *in silico* data shows that compared to all other modification sites K51 is most sterically impaired by acetylation. Longer sidechain modifications at K51 such as succinylation will most likely prevent EfpL from binding to the ribosome.

**Fig. 4:**
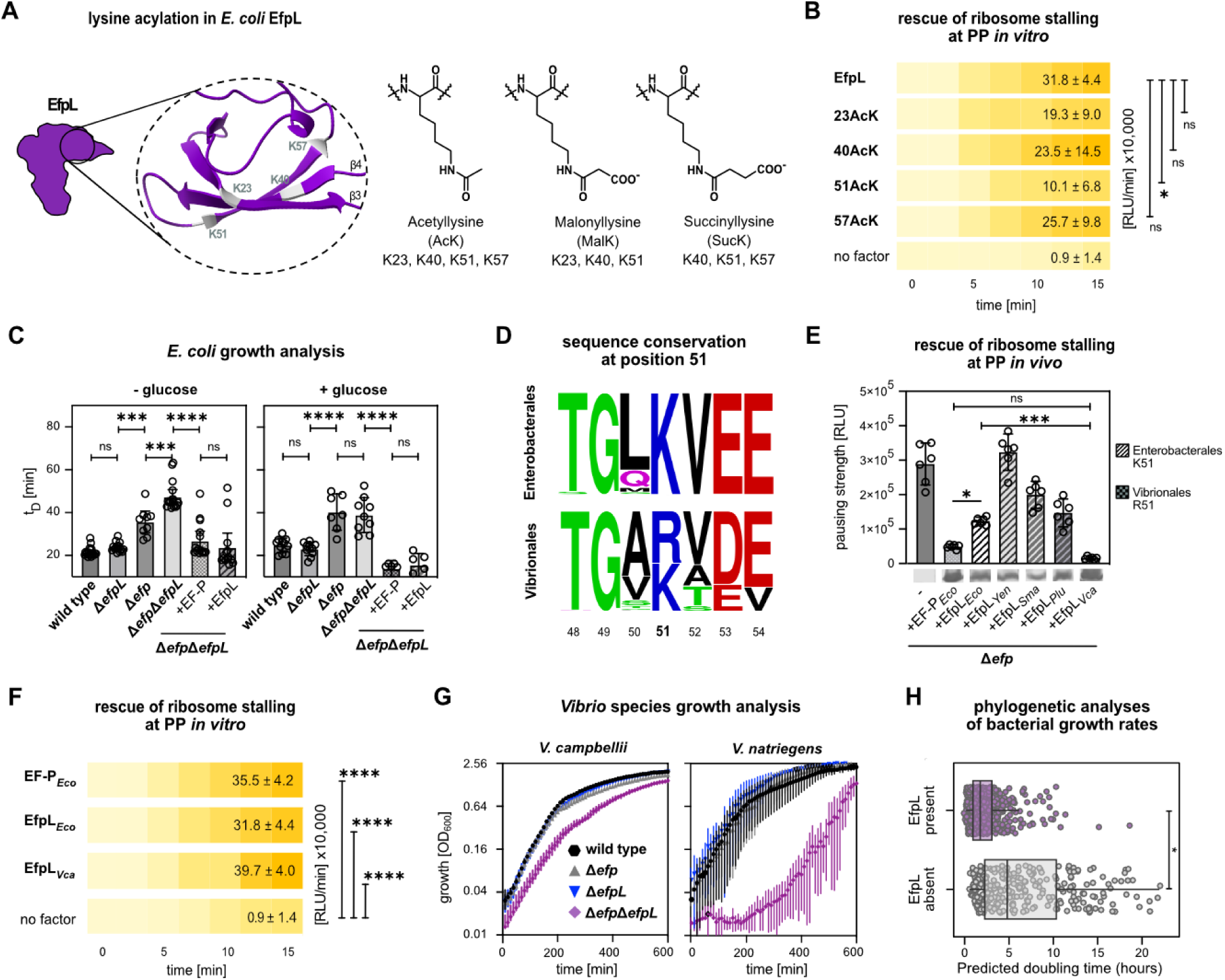
EfpL acylation and its regulation in distinct bacteria. **A)** EfpL acylations according to (Kuhn *et al.*, 2014, Weinert *et al.*, 2013b, Qian *et al.*, 2016). **(B)** *In vitro* transcription and translation of the *nLuc®* variant nLuc_PPN. The absence (no factor) or presence of the respective translation elongation factors of *E. coli* EF-P and EfpL as well as the corresponding substitution variants EfpL_K23AcK, EfpL_K40AcK, EfpL_51AcK, EfpL_K57AcK is shown. Translational output was determined by measuring bioluminescence in a time course of 15 minutes and is given in relative light units measured at the end of the reaction (RLU ± standard deviation) (n ≥ 3). Statistically significant differences according to ordinary one-way ANOVA test (*P value <0.0332, **P value <0.0021, ***P value <0.0002, ****P value <0.0001, ns not significant). **(C)** Growth analysis of *E. coli* BW25113 wild type and deletion strains in LB with and without addition of 20 mM glucose. For complementation *efp* (+EF-P) or *efpL* (+EfpL) were provided in trans. Doubling times (t_D_) were calculated from exponentially grown cells in LB (n ≥ 5). (Statistics and labeling as in (B)). **(D)** Sequence logos^25^ of position 51 and ± 3 amino acids in EfpL in Enterobacterales and Vibrionales. **(E)** *In vivo* comparison of pausing at PPN in *E. coli* Δ*efp* cells and respective trans complementations with *E. coli* EF-P (+EF-P*_Eco_*), *Yersinia enterocolitica* (+EfpL*_Yen_*), *Serratia marcenscens* (+EfpL*_Sma_*), *P. luminescens* (+EfpL*_Plu_*), as well as *Vibrio campbellii* (+EfpL*_Vca_*). Pausing strength is given in relative light units (RLU) (n=6, Error bars depict standard deviation). (Statistics as in (B)) **(F)** *In vitro* transcription and translation as in (B). The absence (no factor) or presence of the respective translation elongation factors of *E. coli* (EF-P, EfpL) and *V. campbellii* (EfpL*_Vca_*) is shown. (Statistics as in (B)) **(G)** Growth analysis of *V. campbellii* and *Vibrio natriegens* with corresponding mutants lacking *efp (*Δ*efp)*, *efpL (*Δ*efpL)*, or both genes *(ΔefpΔefpL)* in LM and LB, respectively (n = 12; error bars indicate standard deviation). **(H)** Phylogenetic analysis of predicted γ-proteobacterial growth rates comparing EfpLs absence or presence. Doubling times were predicted using codon usage bias in ribosomal proteins (phylogenetic ANOVA, P value = 0.029, n =786, P value is based on 1000 permutations).

Acylation is predominantly a non-enzymatic modification and depends on the metabolic state of a cell^43,44^. Hence, different growth conditions favor or disfavor acylation levels. In this regard glucose utilization promotes acetylation of EfpL due to increased levels of the acetyl group donor acetyl phosphate and is especially pronounced for K51^43,44^. We were therefore curious to compare growth of *E. coli* as well as *efp* and *efpL* mutant strains in LB and LB supplemented with 20 mM of glucose (**Fig. 4C**). Strikingly, when glucose was added to the medium, the previously observed growth phenotype of the Δ*efpL* mutant strains in LB completely disappeared, further corroborating our *in vitro* and *in silico* data of EfpL deactivation by acylation.

It has recently been shown that acetylation of ribosomal proteins in general inhibits translation and increases the proportions of dissociated 30S and 50S ribosomes^48^. In addition to this scheme, we have now uncovered, that in *E. coli* EfpL – but not EF-P – is deactivated by acylation. In this way the EfpL acts as a sensor for the metabolic state to regulate translation of specific XP(P)X proteins.

### EfpL is an evolutionary driver of faster growth

Paralogous proteins evolve to diversify functionality and enable species-specific regulation^49^. In this regard we found that in enterobacteria, the four acylation sites of EfpL in *E. coli* remain largely invariable, whereas in others, such as *Vibrio* species, they show less conservation (**Fig. 4D; Extended Data Fig. 9**). Most importantly, lysine in position 51 is an arginine in the EfpLs of e.g. *Vibrio cholerae, Vibrio natriegens* and *Vibrio campbellii*. Moreover, we found that expression levels of *efpL V. campbellii* (*efpL_Vca_*) are much higher than in *E. coli* and equal those of *efp_Vca_*, together suggesting a broader role for EfpL in this organism (**Supplementary Fig. S9**). We compared the rescue efficiency of EfpL*_Vca_* with those of selected *Enterobacteriacae* (**Fig. 4E**) and found that overproduction of EfpL*_Vca_* was superior over all tested enterobacterial EfpLs. In fact, the protein could most efficiently counteract the translational arrest at PPN not only *in vitro* but also *in vivo* (**Fig. 4E, F; Extended Data Fig. 10**). Next, we investigated the effect of an *efpL_Vca_* deletion. Similar to *E. coli* we did not find any growth phenotype. However, in stark contrast, a deletion of *efp_Vca_* also had no consequences for growth speed. Only the simultaneous deletion of both genes (Δ*efp*Δ*efpL*) diminished growth in *V. campbelli*, suggesting that EfpL*_Vca_* and EF-P*_Vca_* can fully compensate for the absence of the other. To exclude a species specific behavior we further included *V. natriegens,* the world record holder in growth speed (doubling time is less than 10 minutes under optimal conditions) (**Fig. 4G**) ^50^. Similar to *V. campbellii* both proteins seem to be of equal importance. Therefore, we conclude, that the role of EfpL in ribosomal rescue of XP(P)X is more general in *Vibrio* species compared to Enterobacteria.

We were ultimately curious, whether there might be a universal benefit for bacteria in encoding EfpL. To this end, we estimated doubling times of a reference data set of γ-proteobacteria using a codon usage bias-based method (**Extended Data Fig. 11**) ^51^. Then we categorized them according to presence or absence of an EfpL paralog. To minimize differences resulting from phylogenetic diversity we focused specifically on γ-proteobacteria encoding an EF-P that is activated by EpmA (**Fig. 4H**). Strikingly, bacteria with EfpL are predicted to grow faster than those lacking it. Thus, we conclude that the concomitant presence of EF-P and EfpL is an evolutionary driver for faster growth. We speculate that microorganisms with both proteins benefit from their unique capabilities to interact with the P-site tRNA^Pro^, which in turn helps to increase overall translation efficiency.

## Discussion

Proline is the only secondary amino acid in the genetic code. The pyrrolidine ring can equip proteins with unique properties^52^ and the polyproline helix is just one expression for the structural possibilities^53^. However, all this comes at a price. The rigidity of proline decelerates the peptidyl transfer reaction with tRNA^Pro^. Not only is it a poor A-site peptidyl acceptor, but also proline is a poor peptidyl donor for the P-site^54,55^. Nevertheless, arrest inducing polyprolines occur frequently in pro- and eukaryotic genomes^39,56^. This in turn, shows that the benefits of such sequence motifs outweigh the corresponding drawbacks and explain why nature has evolved the universally conserved EF-P to assist in translation elongation at XP(P)X ^30^. To promote binding to the polyproline stalled ribosome EF-P specifically interacts with the D-loop of the P-site tRNA^Pro9^, the L1 stalk, and the 30S subunit^11^ and the mRNA^10^, with the latter being the only variable in this equation. Accordingly, in the ideal case, the EF-P retention time on the ribosome could be modulated according to the motif’s arrest strength. Indeed, the dissociation rate constant of EF-P from the ribosome differs depending on the E-site codon^38^. Our data support the hypothesis that amino acids encoded by a codon beginning with a guanosine induce a particularly strong translational arrest in XP(P) motifs (**Fig. 3E, F; Extended Data Fig. 6; Supplementary Fig. S6**). As EF-P is an ancient translation factor being already present before phylogenetic separation of bacteria and eukaryotes/archaea^57^, we wondered whether there is a connection to the evolution of the genetic code. Remarkably, all six amino acids encoded by GNN (Gly, Ala, Asp, Glu, Val, Leu) are included among the standard amino acids that can be produced under emulated primordial conditions^58^. One might therefore speculate that in the early phase of life, EF-P/IF-5A were essential to assist in nearly every peptide bond formation with proline in the P-site and thus reading the E-site codon by a second OB-domain was especially beneficial.

The importance to alleviate ribosome stalling at consecutive prolines is further underlined by the existence of additional rescue systems namely the <u>A</u>TP-<u>B</u>inding <u>C</u>assette family-<u>F</u> (ABCF) protein Uup in *E. coli* and its ortholog YfmR in *B. subtilis*^37,59–61^. In interplay with EF-P, Uup/YfmR and EfpL can facilitate translation of XP(P)X containing proteins. The different modes of action and structural characteristics of the three factors enabled specialization. In case of EfpL the protein is superior in ribosome rescue at specific genes (**Fig. 3D; Supplementary Tab. S3**). This, in turn, might be an evolutionary driving force for translational speed and hence higher growth rates as indicated by our phylogenetic analysis (**Fig. 4H**). Alternatively, an EF-P paralog opens new regulatory possibilities. In contrast to EfpL encoding bacteria, some lactobacilli, for instance, have two copies of *efp* in the genome (**Supplementary Tab. S1**) ^14^. One might speculate that here one *efp* is constitutively expressed and the second copy is transcriptionally regulated according to the translational needs. Although relying only on one EF-P, such regulation was reported for Actinobacteria, in which polyproline containing proteins are concentrated in the accessory proteome^62^. Here EF-P accumulates during early stationary phase and might boost secondary metabolite production as evidenced for *Streptomyces coelicolor.* By contrast, for *E. coli* EfpL there is no evidence for such copy number control, as it simply mirrors the expression pattern of other ribosomal proteins^28^. Instead, the protein seems to fulfill a dual role in this organism. On the one hand it is essential for full growth speed (**Fig. 2A**; **Fig. 4H**). On the other hand, it acts as sensor of the metabolic state (**Fig. 4C**). The combination of multiple sites of acylation^43–46^ and the chemical diversity of this modification type^63^ lead to a highly heterogenic EfpL population, which could fine tune translation in each cell differently. We speculate that regulation of translation by acylation^48^ in general and of EfpL in particular adds to phenotypic heterogeneity and thus might contribute to survival of a population under changing environmental conditions^64^. Such a scenario is particularly important for bacteria that colonize very different ecological niches, such as many enterobacteria including *E. coli* do. Depending on whether they are found e.g. in the soil/water or in the large intestine, the nutrient sources they rely on change. Therefore, it is plausible to assume that fine-tuning metabolic responses by acylating and deacylating, EfpL gives enterobacteria an advantage to thrive in the gastrointestinal tract.

Compared to the eukaryotic and archaeal IF-5A, EF-P diversity is much greater^8,65^. Especially the functionally significant β3Ωβ4 has undergone significant changes. Starting with the catalytic residue at the loop tip, which is not restricted to lysine as for eukaryotes/archaea. Instead, one also finds asparagine, glutamine, methionine, serine and glycine, besides arginine^23^. These changes extend to the overall sequence composition of β3Ωβ4 to either increase stiffness^66^ or, in the case of EfpL, to prolong the loop, as shown in this study by the first EfpL high-resolution structure. The latter two strategies functionalize the protein without modification. Interestingly, however, the EfpL subgroup is phylogenetically linked most closely to the EF-P branch being activated by α-rhamnosylation^14,21^. This raises the question about the evolutionary origin of EfpL. Starting from a lysine type EF-P^57^, we speculate that upon gene duplication and sequence diversification an early EfpL arose, and cells benefitted from improved functionality in a subset of XP(P)X arrest peptides. Further evolutionary events could include the shrinkage of the loop back to the canonical seven amino acids and eventually the phylogenetic recruitment of EarP. Such phylogenetic order is supported by an invariant proline upstream of the catalytically active loop tip residue which is found in EfpLs and lysine type EF-Ps, but is absent in EarP type EF-Ps (**Fig. 1**).

Lastly, EF-P diversity holds also potential for synthetic biology applications. Reportedly, EF-P can boost peptide bond formation with many non-canonical amino acids (ncAA) ^67–70^. This includes not only proline derivatives but also D- and β-amino acids. However, in all studies *E. coli* EF-P was used. Given the structural differences between EfpL and EF-P and the resulting differences in the rescue spectrum, we speculate that use of EfpL might be especially beneficial for genetic code expansion for certain ncAA.

Collectively, our structural and functional characterization of the EfpL subfamily not only underscores the importance of ribosome rescue at XP(P)X motifs, but also adds another weapon to the bacterial arsenal for coping with this type of translational stress. We further illustrate how different bacteria utilize this weapon to gain evolutionary advantages and give an outlook on how EfpL can potentially be used as a molecular tool.

## Methods

### Plasmid and strain construction

All strains, plasmids, and oligonucleotides used in this study are listed and described in Supplementary data files (**Supplementary Tab. S4**), respectively. Kits and enzymes were used according to manufacturer’s instructions. Plasmid DNA was isolated using the Zyppy® Plasmid Miniprep Kit from Zymo Research. DNA fragments were purified from agarose gels using the Zymoclean® Gel DNA Recovery Kit or from PCR reactions using the DNA Clean & Concentrator®-5 DNA kit from Zymo Research. All restriction enzymes, DNA modifying enzymes, and the Q5® high fidelity DNA polymerase for PCR amplification were purchased from New England BioLabs.

Plasmids for expression of C-termially His_6_-tagged *efp* and *efpL* genes under the control of an inducible promoter were generated by amplification of the corresponding genes from genomic DNA using specific primers and subsequent cut/ligation into the pBAD33 vector^71^. Plasmids for expression of SUMO-tagged *efpL* genes were generated with the Champion™ pET-SUMO Expression System from Invitrogen™ according to manufacturer’s instructions. HisL*_lux reporter strains were generated according to Krafczyk *et al.* ^30^. Deletions and chromosomal integrations of His_6_-tagged encoding genes using RecA mediated homologous recombination with pNPTS138-R6KT of *efp* and *efpL*, were made according to Lassak *et al.* ^72,73^. Genetic manipulations via Red®/ET® recombination were done with the Quick & Easy *E. coli* Gene Deletion Kit (Gene Bridges, Heidelberg, Germany). Reporter plasmid constructions with pBBR1-MCS5-TT-RBS-lux were made according to Gödeke *et al.* ^74^.

### Growth conditions

*E. coli* cells were routinely grown in Miller modified Lysogeny Broth (LB) ^75,76^, super optimal broth (SOB) ^77^ or M9 minimal medium supplemented with 20 mM of Glucose^75^ at 37 °C aerobically under agitation unless indicated otherwise. *V. campbellii* cells were grown in Luria marine (LM) medium (Lysogeny broth supplemented with an additional 10 g/l NaCl) ^78^ at 30 °C aerobically. *V. natriegens* cells were grown in LB at 30 °C aerobically. Growth was recorded by measuring the optical density at a wavelength of 600 nm (OD_600_). When required 1.5 % (w/v) agar was used to solidify media. Alternative carbon sources and media supplements were added and are indicated. If needed, antibiotics were added at the following concentrations: 100 µg/ml carbenicillin sodium salt, 50 µg/ml kanamycin sulfate, 20 µg/ml gentamycin sulfate, 30 µg/ml chloramphenicol. Plasmids carrying pBAD^71^ or Lac promoter were induced with ւ(+)-arabinose at a final concentration of 0.2 % (w/v) or Isopropyl-β-D-thiogalactopyranosid (IPTG) at a final concentration of 1 mM, respectively.

### *In vivo* promotor activity assay

*E. coli* cells harboring the plasmids pBBR1-MCS5-P*_efp_*-*luxCDABE* or pBBR1-MCS5-P*_efpL_*-*luxCDABE* were inoculated in LB and appropriate antibiotics and were grown aerobically at 37 °C. The next day, 96-well microtiter plates with fresh LB and antibiotics were inoculated with the cells at an OD_600_ of 0.01. The cells were grown aerobically in the CLARIOstar® PLUS at 37 °C. OD_600_ and luminescence were recorded in 10 min intervals over the course of 16 h. Light emission was normalized to OD_600_. Each measurement was performed in triplicates as a minimum.

### LDC Assay

Cells were cultivated in LDC indicator medium (indicator: bromothymol blue) for 16 h and the pH increase was shown qualitatively as a color change^3^.

### MgtL reporter assay

*E. coli* cells harboring the plasmids pBBR1-MCS5-*mgtL_luxCDABE* were inoculated in M9 minimal supplemented with the appropriate antibiotics and grown aerobically at 37 °C. The next day, a microtiter plate with fresh M9 minimal medium initially leaving out Mg^2+^ (Mg^2+^-free M9). Indicated concentrations of Mg^2+^ (added as MgSO_4_) were added subsequently.Cells were inoculated with a starting OD_600_ of 0.01. Then cells were grown aerobically in the CLARIOstar^®^ PLUS at 37 °C. OD_600_ and luminescence was recorded in 10 min intervals over the course of 16 h. Light emission was normalized to OD_600_. Each measurement was performed in triplicates as a minimum.

### Measurement of pausing strength *in vivo*

The pausing strength of different motifs was determined according to Krafczyk *et al.* ^30^ by measuring absorption at 600 nm (Number of flashes: 10; Settle time: 50 ms) and luminescence emission (Attenuation: none; Settle time: 50 ms; Integration time: 200 ms) with a Tecan Infinity® or ClarioStar plate reader in between 10-min cycles of agitation (orbital, 180 rpm, amplitude: 3 mm) for around 16 h.

### Protein overproduction and purification

For *in vitro* studies C-terminally His_6_-tagged EF-P and EfpL variants were overproduced in *E. coli* LMG194 harboring the corresponding pBAD33 plasmid. C-terminally His_6_-tagged EfpL with acetyl lysine instead of lysine at position 23, 40, 51 or 57 were overexpressed from pBAD33_*efpL*K23Amber_His_6_, pBAD33_*efpL*K40Amber_His_6_, pBAD33_*efpL*K51Amber_His_6_, or pBAD33_*efpL*K57Amber_His_6_ in *E. coli* LMG194 which contained the additional plasmid pACycDuet_AcKRST described in Volkwein *et al*. ^47^. This allowed for amber suppression utilizing the acetyl lysine-tRNA synthetase (AcKRS) in conjunction with PylT-tRNA. LB was supplemented with 5 mM *N*^ε^-acetyl-L-lysine and 1 mM nicotinamide to prevent deacetylation by CobB^79^. During exponential growth, 0.2 % (w/v) ւ(+)- arabinose was added to induce gene expression from pBAD vectors, and 1 mm IPTG served to induce gene expression of the pACycDuet-based system. Cells were grown overnight at 18 °C and harvested by centrifugation on the next day. The resulting pellet was resuspended in HEPES buffer (50 mM HEPES, 100 mM NaCl, 50 mM KCl, 10 mM MgCl_2_, 5 % (w/v) glycerol, pH 7.0). Cells were then lysed using a continuous-flow cabinet from Constant Systems Ltd. (Daventry, UK) at 1.35 kbar. The resulting lysates were clarified by centrifugation at 4 °C at 234 998 *g* for 1 h. The His_6_-tagged proteins were purified using Ni-NTA beads (Qiagen, Hilden, Germany) according to the manufacturer’s instructions, using 20 mM imidazole for washing and 250 mM imidazole for elution. In the final step, the purified protein was dialyzed overnight against HEPES buffer to remove imidazole from the eluate. For MS analysis cells with chromosomally encoded His_6_-tagged EfpL were grown in SOB until mid-exponential growth phase and harvested by centrifugation. To overproduce EfpL proteins LMG194 harboring a pBAD33 plasmid with C-terminally His_6_-tagged EfpL were grown in SOB and supplemented with 0.2 % (w/v) ւ(+)-arabinose during exponential growth phase (OD_600_). Cells were grown overnight at 18 °C and harvested by centrifugation on the next day. Pellets were resuspended in 0.1 M sodium phosphate buffer, pH 7.6. Cells were then lysed using a continuous-flow cabinet from Constant Systems Ltd. (Daventry, UK) at 1.35 kbar. The resulting lysates were clarified by centrifugation at 4 °C at 234 998 ***g*** for 1 h. The His_6_-tagged proteins were purified using Ni-NTA beads (Qiagen, Hilden, Germany) according to the manufacturer’s instructions. For washing and elution, a gradient of imidazole (10, 25, 50, 75, 100, 150, 200, 250 mM) was used. The purified protein was dialyzed overnight against in 0.1 M sodium phosphate buffer, pH 7.6 to remove imidazole from the eluate.

For crystallization BL21 cells harboring a pET-SUMO plasmid were grown in SOB and supplemented with 1 mM IPTG during the exponential growth phase. Cells were grown overnight at 18 °C and harvested by centrifugation on the next. Pellets were resuspended in 0.5 M Tris-HCl buffer, pH 7.0. Cells were then lysed using a continuous-flow cabinet from Constant Systems Ltd. (Daventry, UK) at 1.35 kbar. The resulting lysates were clarified by centrifugation at 4 °C at 234 998 ***g*** for 1 h. The His_6_-tagged proteins were purified using Ni-NTA beads (Qiagen, Hilden, Germany) according to the manufacturer’s instructions, using 20 mM imidazole for washing and 250 mM imidazole for elution. The purified protein was dialyzed overnight against 0.5 M Tris-HCl buffer, pH 7.0 to remove imidazole from the eluate. 0.33 mg SUMO-protease per 1 mg protein were added and incubated over night at 4 °C. SUMO-protease and SUMO-tag were captured using Ni-NTA beads (Qiagen, Hilden, Germany) according to the manufacturer’s instructions. The protein was additionally purified via size exclusion chromatography on a Superdex 75 10/300 Increase column (Cytiva) in 20 mM Tris-HCl, 50 mM NaCl and 1 mM DTT at pH 8.0. Fractions with the protein of interest were concentrated and further subjected to anion exchange chromatography on a Resource Q (Bio-Rad) 6 ml-column to remove remaining contaminants with a NaCl salt gradient from 50 mM to 500 mM. The protein eluted at approximately 200 mM NaCl. The final sample was buffer-adjusted to 50 mM NaCl for crystallization.

### SDS–PAGE and western blotting

For protein analyses cells were subjected to 12.5 % (w/v) sodium dodecyl sulfate (SDS) polyacrylamide gel electrophoresis (PAGE) as described by Laemmli^80^. To visualize proteins by UV light 2,2,2-trichloroethanol was added to the polyacrylamide gels^81^. Subsequently, the proteins were transferred onto nitrocellulose membranes, which were then subjected to immunoblotting. In a first step the membranes were incubated either with 0.1 μg/ml anti-6×His® antibody (Abcam). This primary antibody, produced in rabbit, were targeted with 0.2 μg/ml anti-rabbit alkaline phosphatase-conjugated secondary antibody (Rockland) or 0.1 µg/ml anti-rabbit IgG (IRDye® 680RD) (donkey) antibodies (Abcam). Anti-rabbit alkaline phosphatase-conjugated secondary antibody was detected by adding development solution [50 mM sodium carbonate buffer, pH 9.5, 0.01 % (w/v) p-nitro blue tetrazolium chloride (NBT), and 0.045 % (w/v) 5-bromo-4-chloro-3-indolyl-phosphate (BCIP)]. Anti-rabbit IgG were visualized via Odyssey® CLx Imaging System (LI-COR, Inc).

### *In vitro* transcription/translation assay

The PURExpress In Vitro Protein Synthesis Kit from New England Biolabs was used according to the manufacturer’s instructions, but reactions were supplemented with EF-P or EfpL, respectively, and a plasmid coding for *nluc* variants (**Supplementary Tab. S4**). Luminescence was measured over time. For a 12.5 μl reaction mixture, 5 μl of PURExpress solution A and 3.75 μl of solution B, 0.25 μl of Murine RNAse inhibitor (New England Biolabs), 5 μM EF-P or EfpL, and 1 ng pET16b_nluc variants are incubated under agitation (300 rpm) at 37 °C. At various time points, a 1 μl aliquot was quenched with 1 μl of 50 mg/ml kanamycin and stored on ice. Afterward, 2 μl of Nano-Glo Luciferase Assay Reagent (Promega) and 18 μl ddH_2_O were added to induce luminescence development, which was detected by the Infinite F500 microplate reader (Tecan®). At least three independent replicates were analyzed, and the statistical significance of the result was determined using GraphPad prism.

### Ribosome Profiling

*E. coli* strains BW25113, BW25113 Δ*efpL*, BW25113 Δ*efp* and BW25113 Δ*efp* complemented with pBAD33-*efpL*_His6 (+EfpL) were cultivated in LB or LB supplemented with 30 µg/mL chloramphenicol and 0.2% ւ-(+)-arabinose at 37 °C under aerobic conditions. Stranded mRNA-seq and ribosome profiling (Ribo-seq) libraries were generated by EIRNA Bio (https://eirnabio.com) from stab cultures. *E. coli* strains were grown in 400 mL LB at 37 °C to an OD600 of 0.4. Cells were harvested from 200 mL of culture by rapid filtration through a Kontes 90mm filtration apparatus with 0.45 μm nitrocellulose filters (Whatman). Cells were scraped from the filter in two aliquots (90% for Ribo-seq / 10% for RNA-seq) before being immediately frozen in liquid nitrogen. Total RNA was extracted from RNA-seq aliquots in trizol before mRNA was rRNA depleted, fractionated, and converted into Illumina compatible cDNA libraries. Ribo-seq aliquots were lysed in 600 μl ice-cold polysome lysis buffer (20 mM Tris pH 8; 150 mM MgCl_2_; 100 mM NH_4_Cl; 5 mM CaCl_2_; 0.4 % Triton X-100; 0.1 % NP-40; 20 U/ml Superase*In; 25U/mM Turbo DNase) by beadbeating in a FastPrep-24 with CoolPrep Adapter - 3 rounds at 6 m/s for 30 secs in 2mL cyrovials containing 0.1 mm silica beads. Lysates were clarified by centrifugation at 10,000 g for 5 minutes at 4 °C. Ribosomes for subsequently pelleted from lysates by ultracentrifugation at 370,000 g for 1 hour at 4 °C and resuspended in polysome digestion buffer (20 mM Tris pH 8; 15 mM MgCl_2_; 100 mM NH_4_Cl; 5 mM CaCl_2_). Samples were then digested with 750 U MNase for 1 hour at 25 °C and the reaction was stopped by adding EGTA to a final concentration of 6 mM. Following RNA purification and size selection of ribosome protected mRNA fragments between 20-40 nt in length on 15 % urea PAGE gels, contaminating rRNA was depleted from samples using EIRNA Bio’s custom biotinylated rRNA depletion oligos for *E. coli* before the enriched fragments were converted into Illumina compatible cDNA libraries.

Both stranded mRNA-seq libraries and Ribo-seq libraries were sequenced in three replicates on Illumina’s Nova-seq 6000 platform in 150PE mode to depths of 10 million and 30 million raw read pairs per sample respectively.

The sequence structure of the Riboseq reads was as follows:

QQQ - rpf sequence - NNNNN - BBBBB - AGATCGGAAGAGCACACGTCTGAA,

where Q = Untemplated Addition, rpf sequence = the sequence of the read, N = UMI, a 5 nt are unique molecular identifiers (UMIs), B = Barcode, used to demultiplex (the fastq files have already been demultiplexed) and AGATCGGAAGAGCACACGTCTGAA is the sequence of the adapter. Cutadapt^82^ was used with parameters -u 3 and -a AGATCGGAAGAGCACACGTCTGAA to remove untemplated addition and linker sequence. Untrimmed reads and those shorter than 30 nt after trimming were discarded. Next, the UMI and Barcode was removed and the UMI was used to remove duplicate sequences using a custom python script. Both the Ribo-seq and RNA-seq reads were next mapped to rRNA and tRNA sequences using Bowtie version 1.2^83^. Five Ribo-seq samples were sequenced with two sequencing runs. These samples (WT_Rep3, DELTAefpL_Rep2, DELTAefpL_Rep3, DELTAefp_Rep3 and DELTAefp_plus_efpL_Rep1) were concatenated at this stage. Next the reads were aligned to BW25113 *E. coli* genome (RefSeq accession number NZ_CP009273.1) with Bowtie using parameters (-m 1 -l 25 - n 2 -S). BAM file containing read alignments are available at the SRA archive (ID PRJNA1092679).

The A-site offset in the Riboseq reads was estimated to be 11 nucleotides upstream of the 3’ of the mapped reads. For both Riboseq and RNAseq reads this “A-site” position was used to indicate the genomic location of reads. Pause prediction was carried out on all Ribo-seq samples using PausePred^31^ with a minimum fold-change for a pause score set at 20 within two sliding window sizes of 1000 nt with a minimum coverage of 5 % in the window. The analysis was carried out on aggregated alignment files that included all replicates for each strain. The frequencies of occurrence of trimers of amino acid residues at the locations identified to be pauses were calculated for all possible trimers of amino acid residues. For each trimer of amino acid residues its frequency to be covered by the ribosome in the pause sites was calculated and normalized by dividing by the averaged frequency of the corresponding trimer to occur in the whole ribosome-protected fragments.

### Sequence Data and Domain Analysis

HMMER v.3.4 was used to search for Pfam^84^ domains “EFP_N” (KOW-like domain, PF08207.12), “EFP” (OB domain, PF01132.20), and “Elong-fact-P_C” (C-terminal, PF09285.11) in the protein sequences of 5257 complete representative or reference bacterial genomes (RefSeq) ^24^. We identified 5448 proteins from 4736 genome assemblies that contained all three domains mentioned above (e-value cutoff 0.001) and no other PFAM domains. Sequences of “EFP_N” domains from these proteins were multiply aligned using Clustal Omega v.1.2.4^85^ with all default parameters, shown in a multiple sequence alignment (MSA) (**Supplementary Data S1**), and a phylogenetic tree was inferred by FastTree 2^86^, also with all default parameters. The phylogenetic tree in Newick format is available in the Supplementary Materials (**Supplementary Data S2**). The MSA region comprising positions 40-52 corresponds to the β3Ωβ4 loop region KPGKGQA of the EF-P protein from *E. coli* str. K-12 substr. MG1655 (accession number NP_418571.1) ^22^. The sequence of the EfpL protein (NP_416676.4) from *E. coli* str. K-12 substr. MG1655 has an extended β3Ωβ4 loop SPTARGAAT with the R residue at the tip. The phylogenetic tree was annotated according to the length of the β3Ωβ4 loop and the nature of the residue at the tip of the β3Ωβ4 loop. Those 528 sequences that have an extended β3Ωβ4 loop of more than 7 residues and R at the tip of it formed one branch in the phylogenetic tree. Among the sequences belonging to this branch 474 are annotated as “EfpL” or “YeiP” (synonym of EfpL) proteins in the RefSeq database and no other sequences from the list (**Supplementary Tab. S1**) have this annotation. Sequences with an extended β3Ωβ4 loop of more than 7 residues and the R residue at the tip of it are referred to as EfpL. The remaining 4920 sequences constituted the set of EF-P sequences. The dataset covers 4777 genomes: 4111 of them contain only one sequence with the three domains mentioned above, 660 genomes contain two such sequences, and 6 genomes – three such sequences (**Supplementary Tab. S1**). In a separate analysis step Clustal Omega v.1.2.4^85^ and FastTree 2^86^, both with default parameters, were used to multiply align the sequences of KOW-like domains of the EfpL and EF-P proteins from the EfpL-containing genomes (**Supplementary Data S1**) and to build a phylogenetic tree (**Fig. 1A**). We used the ggtree R package^87^ to visualize the phylogenetic trees and annotate them. Sequence logos were built using Weblogo^25^.

EF-P-containing genomes were scanned for the EpmA, EarP and YmfI proteins. EpmA and EarP proteins were defined as single-domain proteins containing the “tRNA-synt 2” (PF00152.20) and “EarP” (PF10093.9) ^14^ domains, respectively. Using HMMER v.3.4 searches we identified these proteins in 1230 and 565 genomes, respectively. Orthologs of the YmfI protein (Uniprot ID: O31767) from *Bacillus subtilis*^20,65^ were obtained using the procedure described in Brewer and Wagner^23^.

### EfpL structure determination

Initial crystallization trials were performed in 96-well SWISSCI plates at a protein concentration of 4.8 mg/ml using the C3 ShotGun (SG1) crystallization screen (Molecular Dimension). Rod-shaped crystals grew after 7 days at 293K. Diffracting crystals were obtained in 100 mM Sodium-HEPES, 20 % (w/v) PEG 8000 and 10 mM Hexaamminecobalt (III) chloride conditions. The crystals were cryoprotected in mother liquor supplemented with 20 % (v/v) glycerol and snap-frozen at 100K. Datasets from cryo-cooled crystals were collected at EMBL P13 beamlines at the PETRA III storage ring of the DESY synchrotron^88^. The crystals belonged to space group P 1 21 1, with unit cell dimensions of a=60.71, b=53.46, and c=64.95 Å. Preprocessed unmerged datasets from autoproc+STARANISO^89^ were further processed in CCP4cloud^90^. Phases were obtained from molecular replacement using the AlphaFold2 model^91,92^ deposited under ID AF-P0A6N8-F1. The structure was built using the automatic model building pipeline ModelCraft^93^, optimized using PDB-REDO^94^, refined in REFMAC5^95^ with manual corrections in Coot^96^. The quality of the built model was validated with the MolProbity server^97^. The final model was visualized in PyMOL version 2.55 (Delano Scientific). The diffraction data collection and refinement statistics are shown in Supplementary Tab. 2 (**Supplementary Tab. S2A; Supplementary Data S3, 4**).

### Docking and modelling of EF-P and EfpL complexes

For the comparative analysis of EF-P with the E-site codons CCG or GCG through a loop in its C-terminal OB-domain we used the available PDB entry 6ENU^10^ as a starting structure. In accordance with prior definitions by the authors, we directly analyzed and visualized available contacts for the EF-P d3 loop1 around the conserved motif _144_GDT_146_ with the present _-3_CCG_-1_ trinucleotide of the peptidyl-tRNA^Pro^. For contacts with a putative GCG, we initially replaced the initial C nucleotide by G in silico using PyMol (Delano Scientific) and monitored the novel contacts using the implemented tools. For a more thorough analysis, we extracted both the GCG trinucleotide and EF-P from the structure and used the two components for an *in silico* docking followed by energy minimization using HADDOCK^27^. Here, we defined protein residues 146, 147 and 151 as active granting full flexibility to the structure and using automated secondary structure recognition and retainment. RNA residue G-3 was defined as active to enable seed contacts. From a total of 116 structures used by for clustering by HADDOCK 49 were found in the best-scoring cluster 1 (**Supplementary Tab. S2B**). Because of very low remaining restraint violation energies, we integrated the best four models to create an average structure used to analyze contacts between EF-P and RNA.

To analyze and compare interactions of EfpL and EF-P KOW domains with the P-site codon CCA through the β3Ωβ4 loop we looked at the available contacts of the loop as given in the PDB entry 6ENU^10^. For a model of EfpL with the trinucleotide, we aligned the EfpL KOW domain as found in our crystal structure with EF-P from PDB entry 6ENU^10^. We extracted the _74_CCA_76_trinucleotide from the latter and used the two components as starting structures for a docking and energy minimization procedure as described above. Nucleotides 74 and 75 were defined as active, and KOW domain residues 30-35 were set as fully flexible with R33 defined as explicitly active. 198 out of the 200 structures provided by HADDOCK were found in the same cluster with no measurable violations (**Supplementary Tab. S2B**).

For all HADDOCK runs, we implemented the following settings and restraints in context of the spatial and energetic constraints of the natural ribosome environment: Protein N- and C-termini were kept uncharged and no phosphates were left at nucleic acid termini. No particular RNA structure restraints have been applied and only polar hydrogens were installed in both components. For the 0^th^ iteration, components were kept at their original positions for an initial energy-minimizing docking step. No random exclusion of ambiguous restraints was included during docking. Passive residues were defined automatically from the non-active ones using a surface distance threshold of 6.5 Å. We used a minimum percentage of relative solvent accessibility of 15 to consider a residue as accessible. In all runs 1000 initial structures were used in rigid body docking over five trials (excluding 180°-rotations of the ligand), from which the best 200 were subjected to an energy minimization step including short molecular dynamics simulations in explicit water. Default settings were used in advanced sampling parameters of the it1 and final solvated steps (Kyte-Doolittle), respectively. Standard HADDOCK settings were applied for clustering of the 200 final structures with a minimum cluster size of 4.

For the *in silico* analysis of modified lysines, respective sidechans were acetylated based on the EfpL crystal structure using PyMol with no further adjustments of rotamers. The modified KOW domain was then structurally aligned with EF-P in PDB entry 6ENU^10^.

### Mass spectrometry for identification of modification status

For top-down EfpL measurements the proteins were desalted on the ZipTip with C4 resin (Millipore, ZTC04S096) and eluted with 50 % (v/v) acetonitrile 0.1 % (v/v) formic acid (FA) buffer resulting in ∼10 μM final protein concentration in 200–400 μl total volume. MS measurements were performed on an Orbitrap Eclipse Tribrid Mass Spectrometer (Thermo Fisher Scientific) via direct injection, a HESI-Spray source (Thermo Fisher Scientific) and FAIMS interface (Thermo Fisher Scientific) in a positive, peptide mode. Typically, the FAIMS compensation voltage (CV) was optimized by a continuous scan. The most intense signal was usually obtained at -7 CV. The MS spectra were acquired with at least 120,000 FWHM, AGC target 100 and 2-5 microscans. The spectra were deconvoluted in Freestyle (Thermo) using the Xtract Deconvolution algorithm.

### Predicted growth rates

We used a set of 871 genomes from the class gammaproteobacteria from the Integrated Microbial Genomes (IMG) database^98^. These genomes were selected to maximize diversity by including only one genome per Average Nucleotide Identity (ANI) cluster. We used CheckM^99^ v1.0.12 to assess the quality of each genome and retained only those that were predicted to be at least 90 % complete and contain less than 5 % contamination. We re-assigned taxonomy using the Genome Taxonomy Database and GTDB-Tool kit (GTDB-Tk) ^100^ version 0.2.2 and removed genomes where the user-reported species did not agree with GTDB (removed 2 genomes). For example, we removed a genome with a user-reported species of *Serratia marcescens 1822* which was sorted to the genus *Rouxiella* by GTDB-Tk. We also removed 14 genomes of endosymbionts from consideration, mainly from the genus *Buchnera*.

We further subset for only those genomes which contained both genes for *epmA* and *epmB* (removed 62 genomes), contained at least one *efp* gene (removed 2 genomes) and had predicted doubling times under 24 hours (removed 15 genomes). This left 786 genomes for our analysis. We identified the genes for epmA, epmB, efp, and efpL (yeiP) using a combination of different functional databases. We identified *epmA* and *epmB* by searching for the COG^101^ function ids COG2269 and COG1509, respectively. We identified *efp* by searching for the Pfam^102^ domain pfam01132. We identified the gene for *efpL* (*yeiP*) by searching for the TIGRfam^103^ annotation TIGR02178. Next, we estimated the doubling time associated with each remaining genome using the R package gRodon^51^ version 1.8.0. gRodon estimates doubling times using codon usage bias in ribosomal proteins. We used phylogenetic ANOVAs to test differences in predicted doubling times between genomes that encode EfpL and those that don’t. Specifically, we used the phylANOVA function from the R package phytools^104^ version 2.0.3, with p-values based on 1000 permutations.. We made the phylogenetic tree required for this function using 43 concatenated conserved marker genes generated by CheckM. We aligned these sequences using MUSCLE^105^ v3.8.1551 and built the phylogenetic tree using IQ-TREE^106^ v1.6.12. We used the model finder feature^107^ included in IQ-TREE to determine the best-fit substitution model for our tree (which was the LG+R10 model). For this section, we performed all statistical analyses and plotting in R version 4.3.2 and created plots using ggplot2^108^ version 3.4.4..

## Acknowledgment

We thank Kirsten Jung, Kerstin Lassak and Wolfram Volkwein for fruitful discussions and constructive criticism. We also thank Giovanni Gallo, Anja Michl, Nina Kim Hartmann, Bhavna Menon, Irem Niran Cagil and Sabine Peschek for technical assistance. Moreover, Jürgen Lassak is grateful for DFG grant LA 3658/5-1. This work was supported by Liebig fellowship from VCI to Pavel Kielkowski. Fei Qi is supported by the National Natural Science Foundation of China (grant No. 32000462). Andreas. Schlundt acknowledges support through funds SCHL2062/2-1 and 2-2 from the German Research Council (DFG) and the Johanna Quandt Young Academy at Goethe (grant number 2019/AS01). Access to synchrotron beamtime at DESY Hamburg was enabled by the Block Allocation Group grant MX939.

## Author contributions

A.Sieber, R.K., A.Schäpers and J.L. constructed strains and plasmids. A.Sieber and R.K. performed the biochemical in *vivo*/*in vitro* characterization of EfpL. A.Schäpers performed qPCR experiments. A.Sieber purified all proteins used for *in vitro* assays, MS analysis and X-ray crystallography. MS experiments and analysis was done by P.K.. The crystallization screen was set up by J.v.E., K.D. and A.Schlundt, and J.v.E., K.D. and A.Schlundt solved the crystal structure of EfpL*_E. coli_*. A.Schlundt performed *in silico* interaction analyses. All bioinformatic analyses were performed by M.P., T.B., F.Q. and D.F.. Ribosome profiling analyses were done by M.P. and D.F.. M.P. and D.F. performed phylogenetic analyses of the EF-P subgroups and T.B. performed phylogenetic analyses of bacterial growth rates. The study was designed by J.L with contributions from R.K. and D.F.. The manuscript was written by A.Sieber, M.P. and J.L. with contributions from A.Schlundt, T.B. and D.F..

## Data availability

The crystal structure of EfpL*_E. coli_* has been deposited under PDB ID 8S8U and will be accessible upon publication. Ribosome profiling data will be available upon publication at SRD ID PRJNA1092679. R scripts and all files needed to reproduce the analyses on predicted growth rates are available at: https://github.com/tessbrewer/EfpL.

## Extended Data

**Extended Data Fig. 1:**
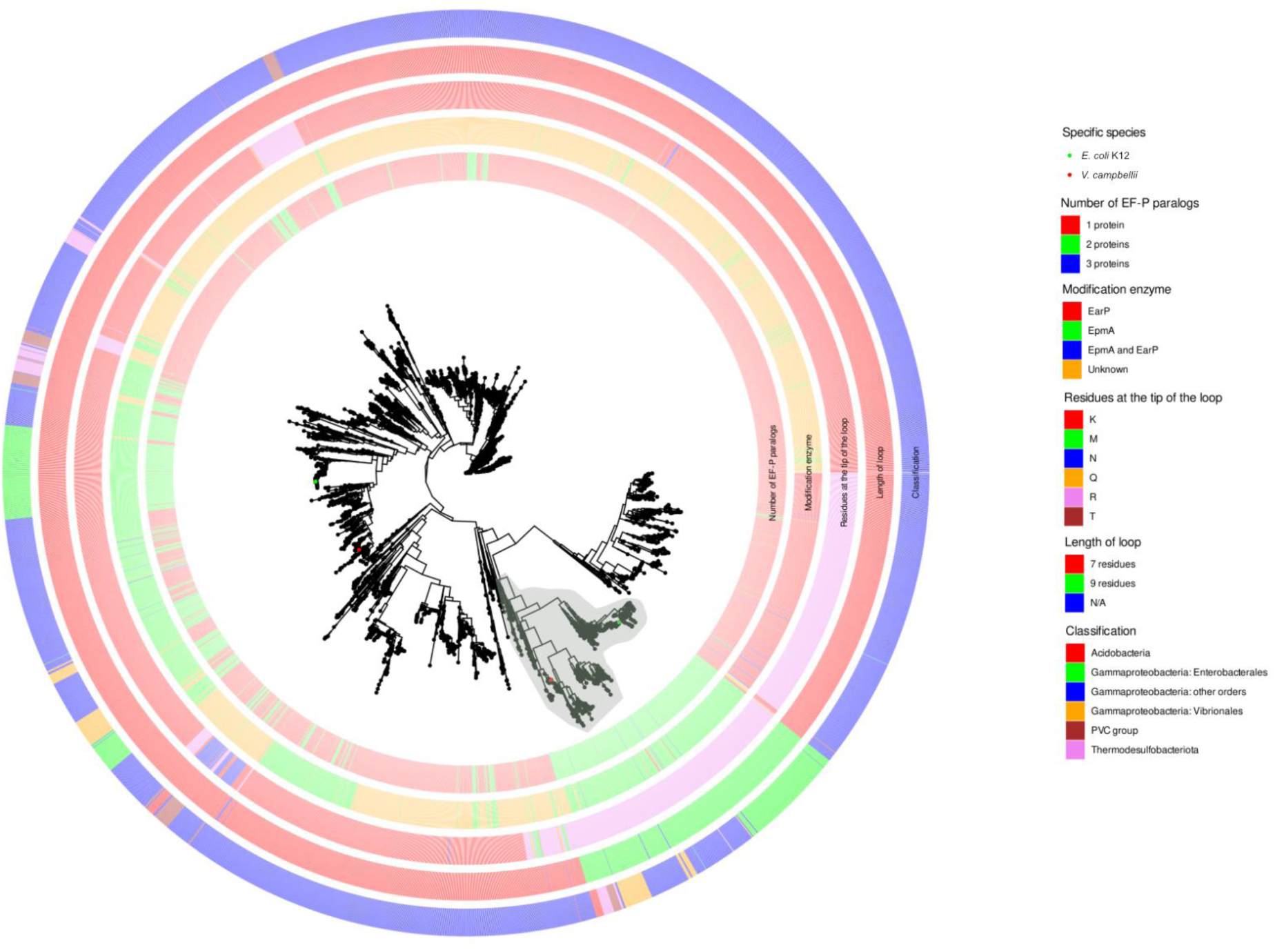
Phylogenetic analysis of EF-P subgroups. Phylogenetic tree was built using the multiply aligned 5448 sequences of KOW-like domains of proteins that have three domains typical for EF-P in a collection of 4736 complete bacterial genomes was obtained from the RefSeq database^24^. Outer ring shows phylogenetic classification in bacterial phyla. Other rings show tip residues and length of β3Ωβ4 loop, as well as number of EF-P homologs and modification enzymes found in bacterial proteomes. Branch endings indicate affiliation to specific species. The green highlighting indicates the branch with protein sequences further annotated as EfpL proteins.

**Extended Data Fig. 2:**
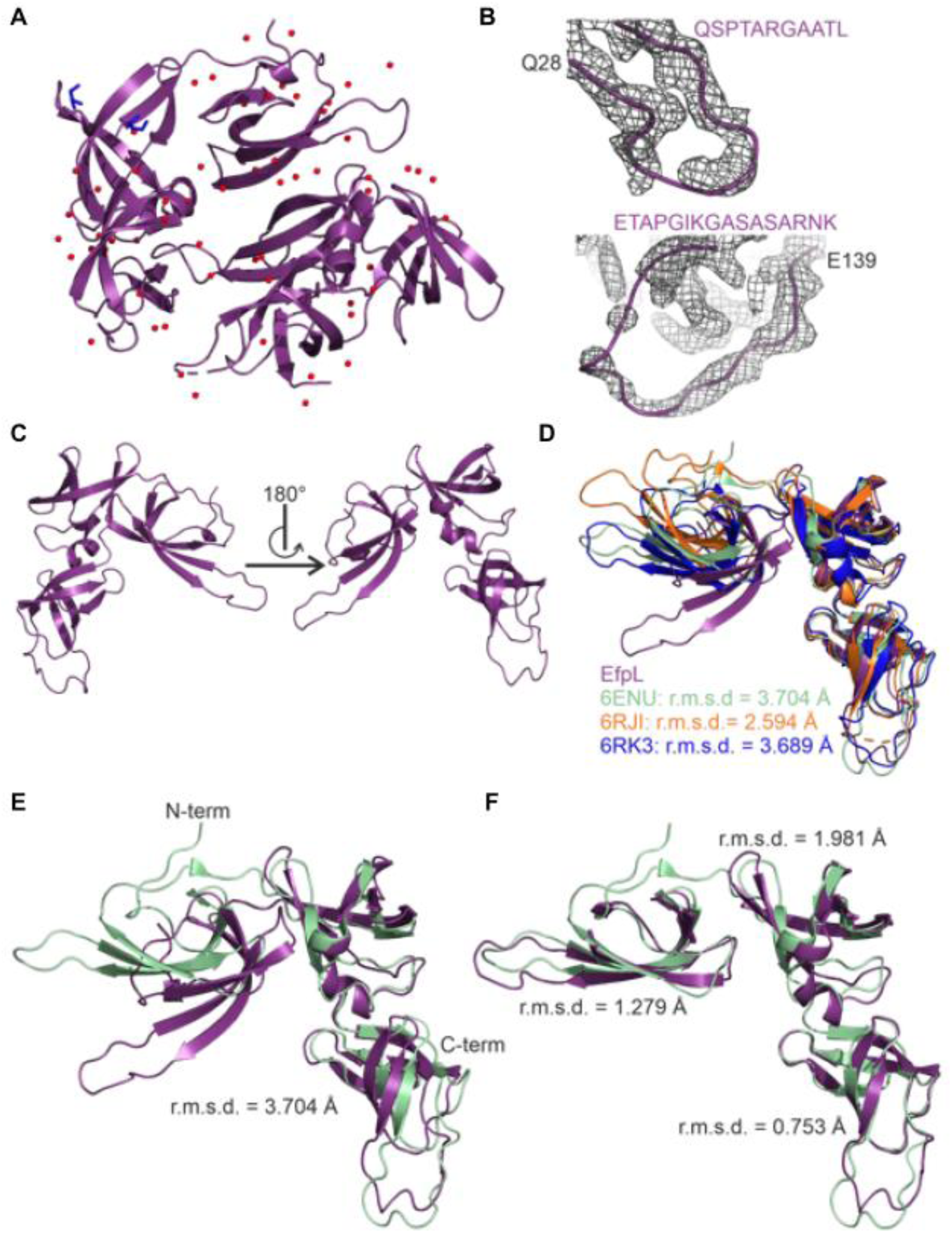
Structural comparison of EfpL and EF-P. **(A)** Structure of crystallographic dimer of EfpL determined by x-ray crystallography in this study. Waters and co-crystallized glycerol ligands are colored in red and blue respectively. **(B)** Electron density map of KOW β3Ωβ4 loop (upper panel) and OB d3 loop 1 (lower panel). The 2Fo-Fc electron density is contoured at 1.5 σ. **(C)** Two-sided view of fully build single chain from the EfpL x-ray structure in A. **(D)** Structural alignment of EfpL with EF-P from *E. coli* (cryo-EM structure, PDB entry 6ENU) and *S. aureus* (crystal structure, PDB entry 6RJI and NMR solution structure, PDB entry 6RK3). R.m.s.d. values in comparison to EfpL are shown. **(E)** Structural alignment of EfpL with EF-P from *E. coli* (cryo-EM structure, PDB entry 6ENU). Root mean square deviation (r.m.s.d) of the total alignment is shown. **(F)** The same as in E but with structured domains (residues 4-56, 68-128, 132-187) of EfpL separated and aligned individually to EF-P and are shown with respective r.m.s.d.

**Extended Data Fig. 3:**
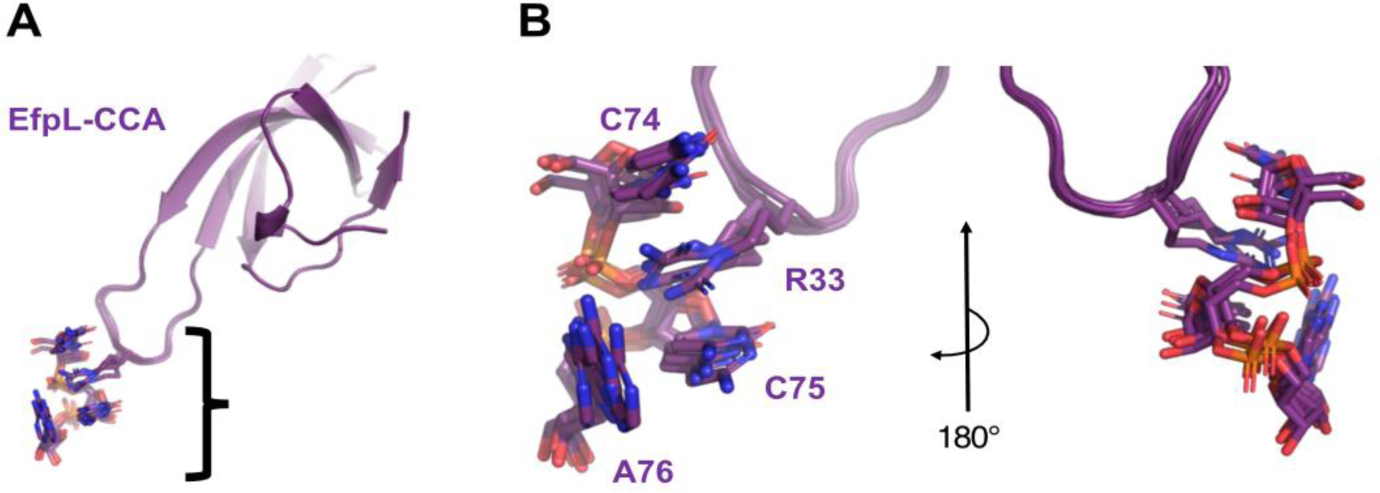
EfpL interaction modeling with the RNA trinucleotide. **(A)** Full-view superimposition of the four best solutions obtained from a HADDOCK run of EfpL together with _74_CCA_76_. **(B)** Zoom-in of panel A to the KOW domain β3Ωβ4 loop region in contact with the RNA trinucleotide. The r.m.s.d. is 0.12 ± 0.01 Å.

**Extended Data Fig. 4:**
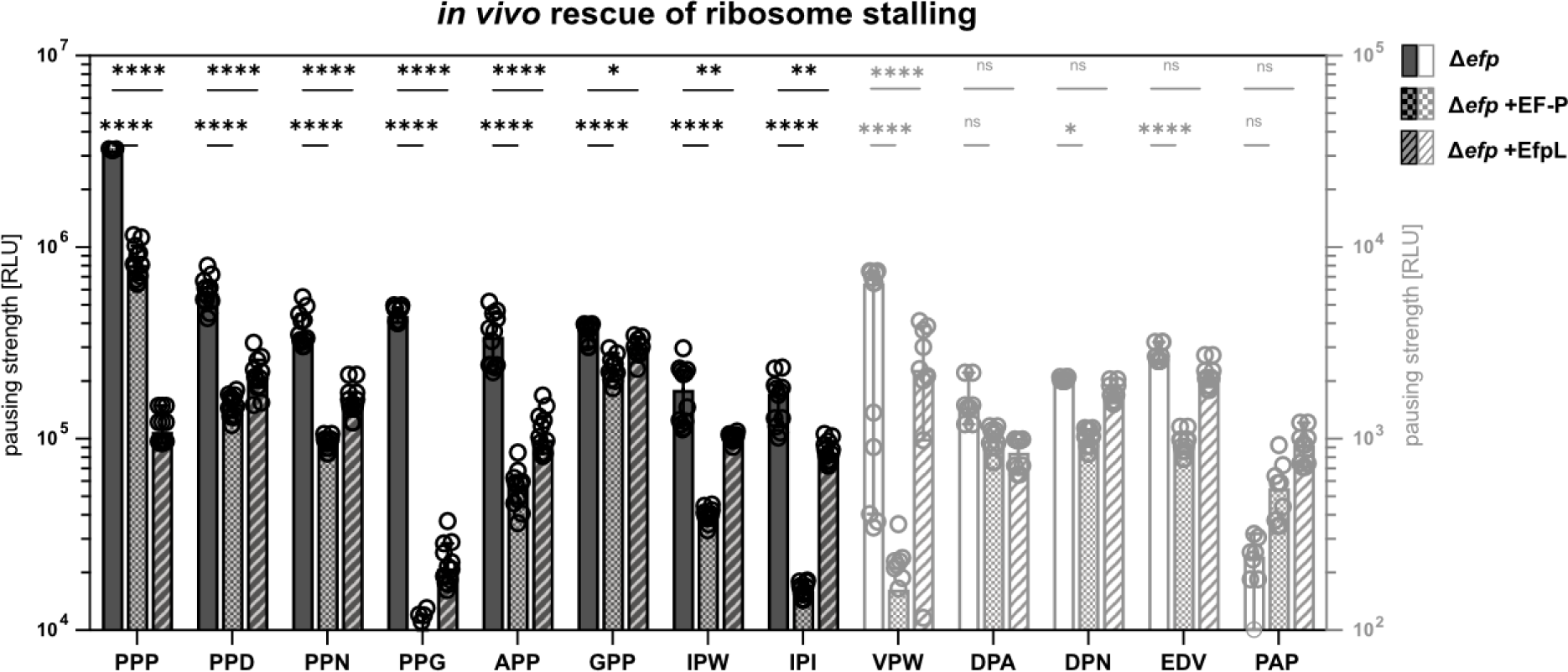
*In vivo* detection of pausing strength at different motifs. *In vivo* comparison of stalling strength of a set of stalling motifs and negative control PAP of *E. coli* Δ*efp* cells and respective trans complementation with EF-P (Δ*efp* +EF-P) and EfpL (Δ*efp* +EfpL). Pausing strength correlates with light emission and is given in relative light units (RLU) (n = 12, Error bars indicate standard deviation). Statistically significant differences according to 2way ANOVA test (*P value <0.0332, **P value <0.0021, ***P value <0.0002, ****P value <0.0001, ns not significant).

**Extended Data Fig. 5:**
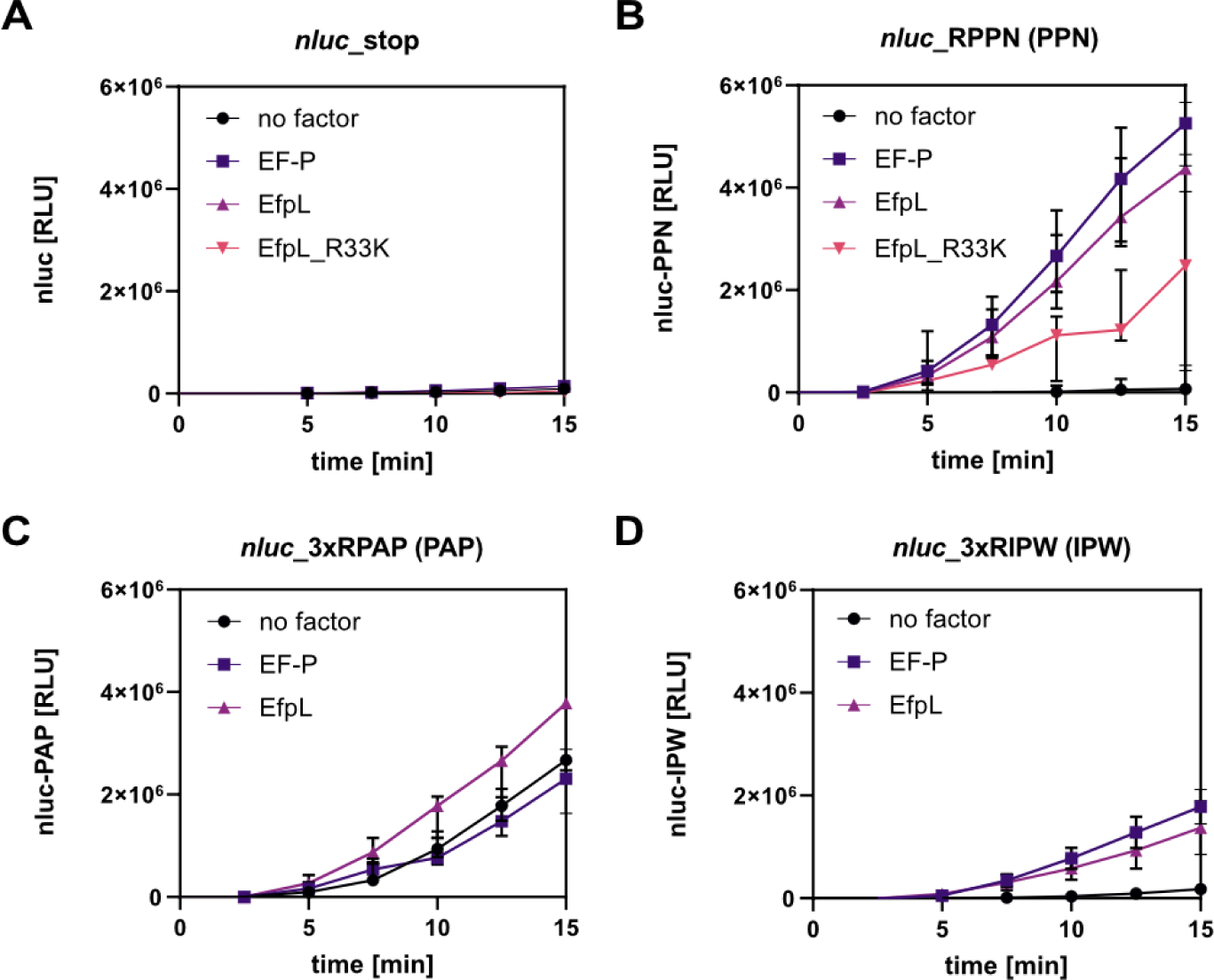
Comparison of EF-P and EfpL of *E. coli* in translating different motifs. *In vitro* transcription and translation of the *nLuc®* variants **(A)** nLuc_stop, **(B)** nLuc_RPPN (PPN), **(C)** nLuc_3xRIPW (IPW) or **(D)** nLuc_3xRPAP (PAP). The absence (no factor) or presence of the respective translation elongation factors of *E. coli* (EF-P, EfpL) is shown. Translational output was determined by measuring bioluminescence in a time course of 15 minutes (n ≥ 3, Error bars indicate standard deviation).

**Extended Data Fig. 6:**
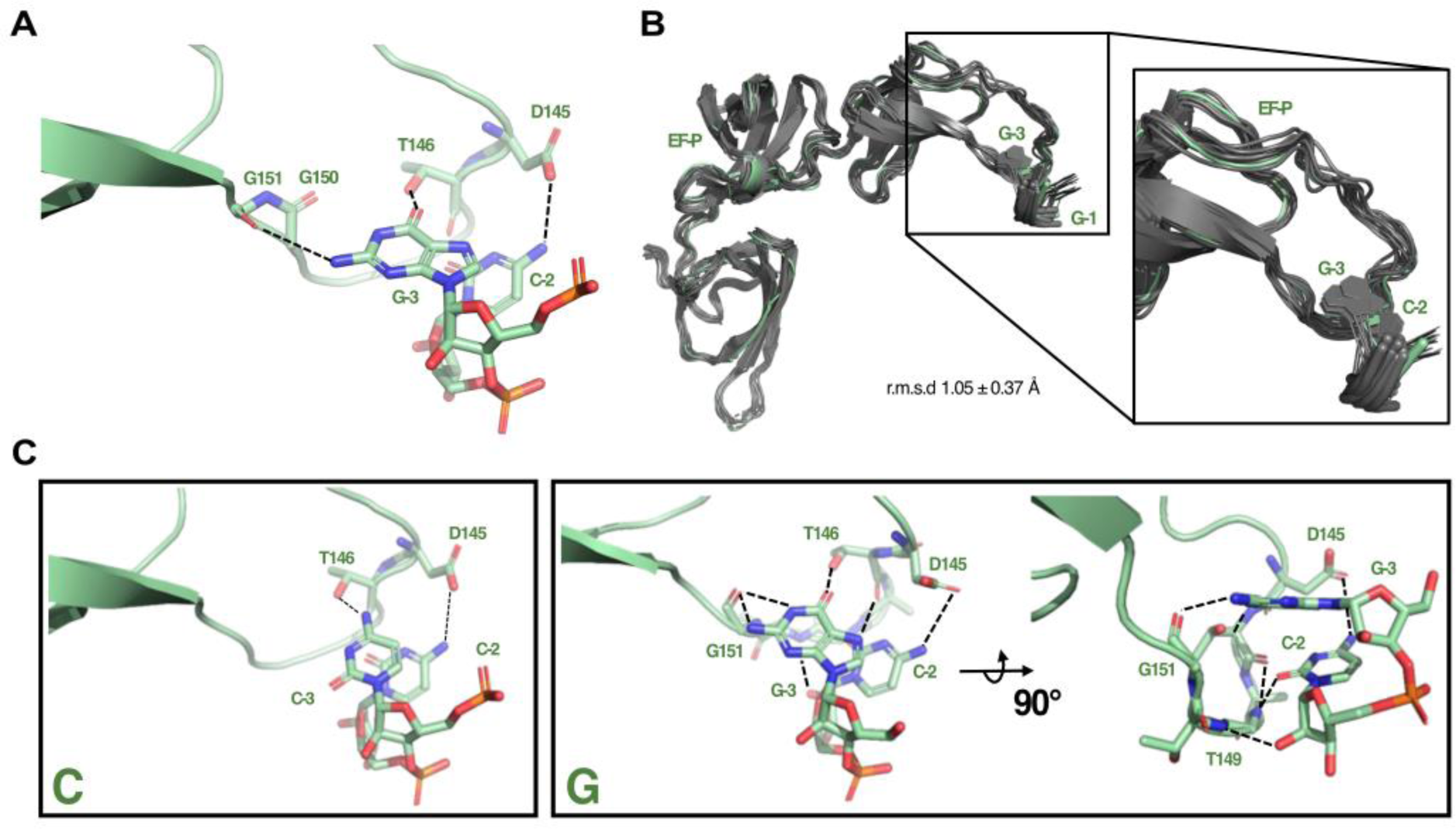
EF-P interaction modelling with E-site codon. **(A)** Close-up view of EF-P in contact with the E-site codon _-3_GCG_-1_, after *in silico* replacement of _-3_C in the PDB entry 6ENU ^10^. Polar contacts to EF-P OB domain 3 loop 1 residues are depicted with broken lines as obtained from the program PyMol (Delano Scientific). Note that only the first two nucleotides are shown for clarity. **(B)** Full-view superimposition of the 10 best solutions obtained from a HADDOCK run of EF-P together with _-3_GCG_-1_. The green model represents the non-docked and non-energy-minimized reference from panel A. The r.m.s.d. is given. The boxed view shows a zoom-in to the EF-P-RNA trinucleotide interface. **(C)** The same as shown in main text Fig. 3, but with an additional perspective depicted for EF-P in complex with GCG to highlight additional contacts.

**Extended Data Fig. 7:**
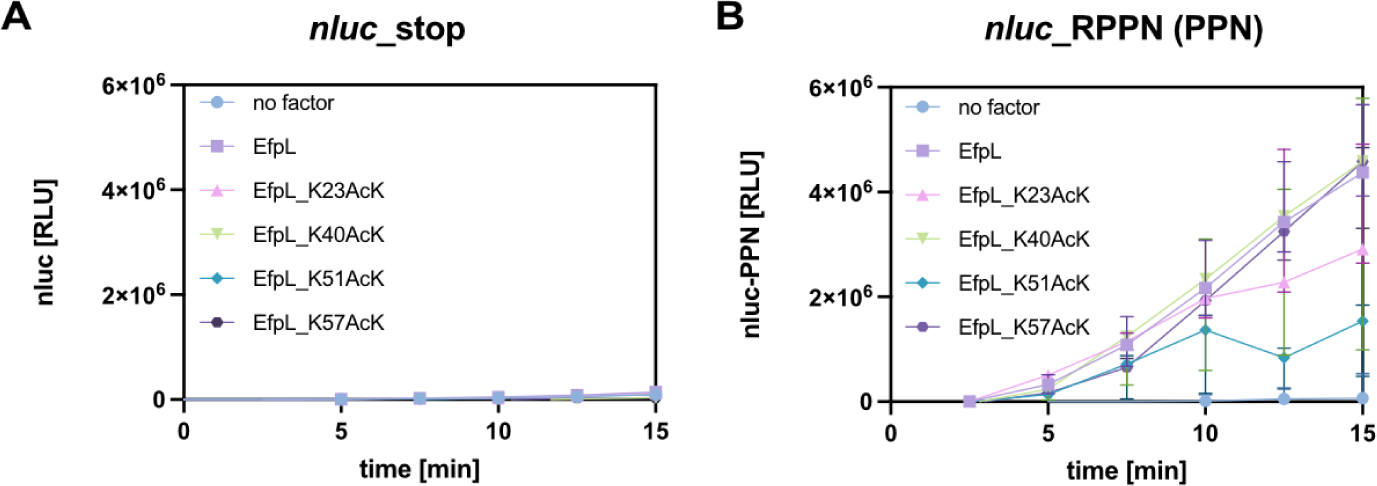
EfpL function dependent on the acylation status. *In vitro* transcription and translation of the *nLuc®* variants **(A)** nLuc_stop (no motif) or **(B)** nLuc_RPPN (PPN). The absence (no factor) or presence of the respective translation elongation factors of *E. coli* EfpL as well as the corresponding substitution variants EfpL_K23AcK, EfpL_K40AcK, EfpL_51AcK, EfpL_K57AcK is shown. Translational output was determined by measuring bioluminescence in a time course of 15 minutes and is given in relative light units (RLU) (n ≥ 3, error bars represent standard deviation). Statistically significant differences according to ordinary one-way ANOVA test (*P value <0.0332, **P value <0.0021, ***P value <0.0002, ****P value <0.0001, ns not significant).

**Extended Data Fig. 8:**
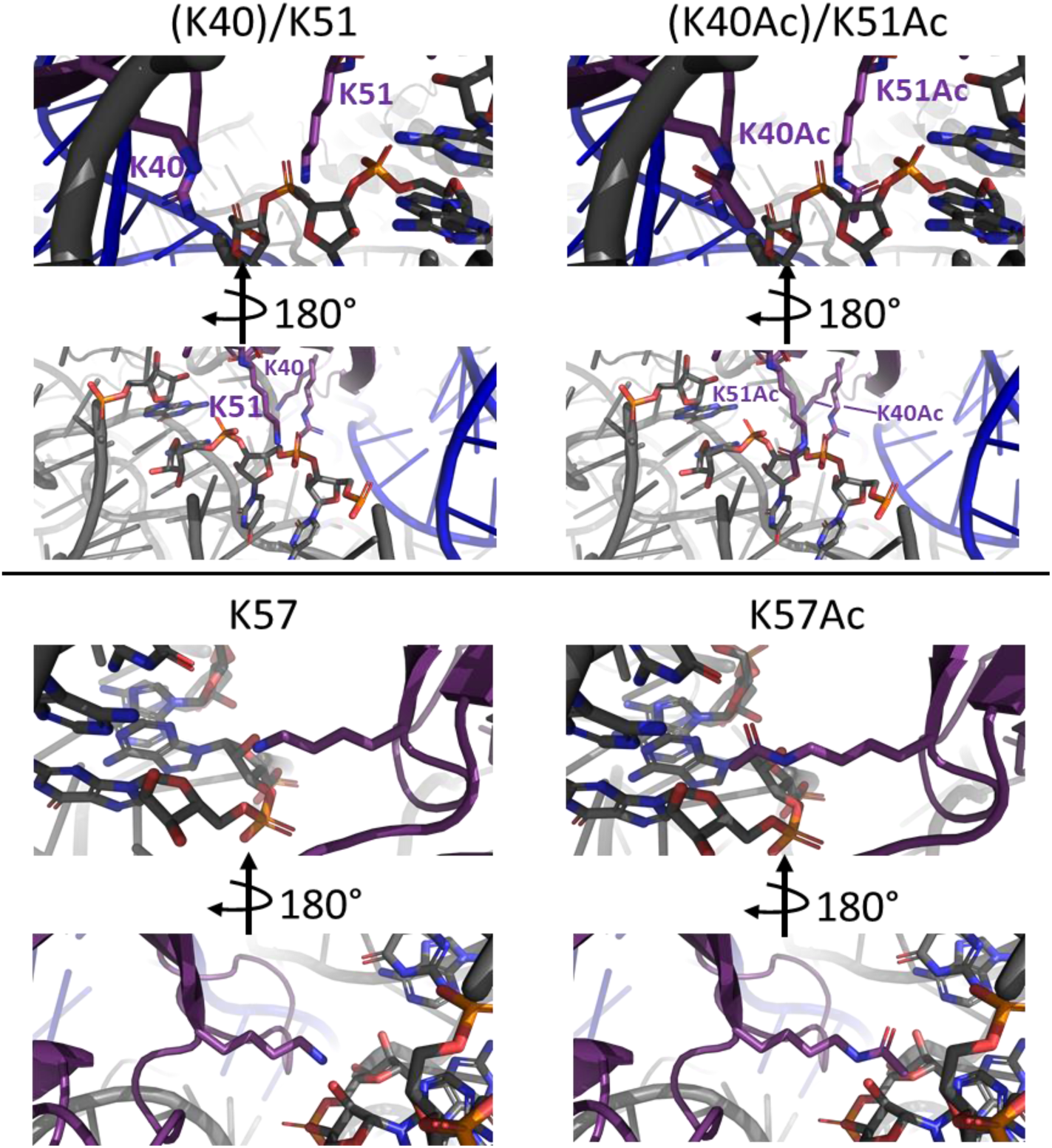
Modelling of Acylation in EfpL. Close-up views on EfpL lysines as shown in unmodified form (left panels) and when acetylated (right panels). Each view is shown from two different perspectives by 180° rotation as indicated. The N-terminal KOW domain (violet-purple) has been aligned to the PDB entry 6ENU^10^ to enable monitoring of potential clashes and interactions with ribosomal components. Lysine sidechains are shown as sticks on an otherwise cartoon-typed presentation. For the K40/K51 region (upper panels), R42 is additionally shown to indicate the dense space, relevant in potential sidechain modifications. Relevant RNA regions in close vicinity of lysines are shown as sticks. Grey represents ribosomal RNA, blue indicates tRNA.

**Extended Data Fig. 9:**
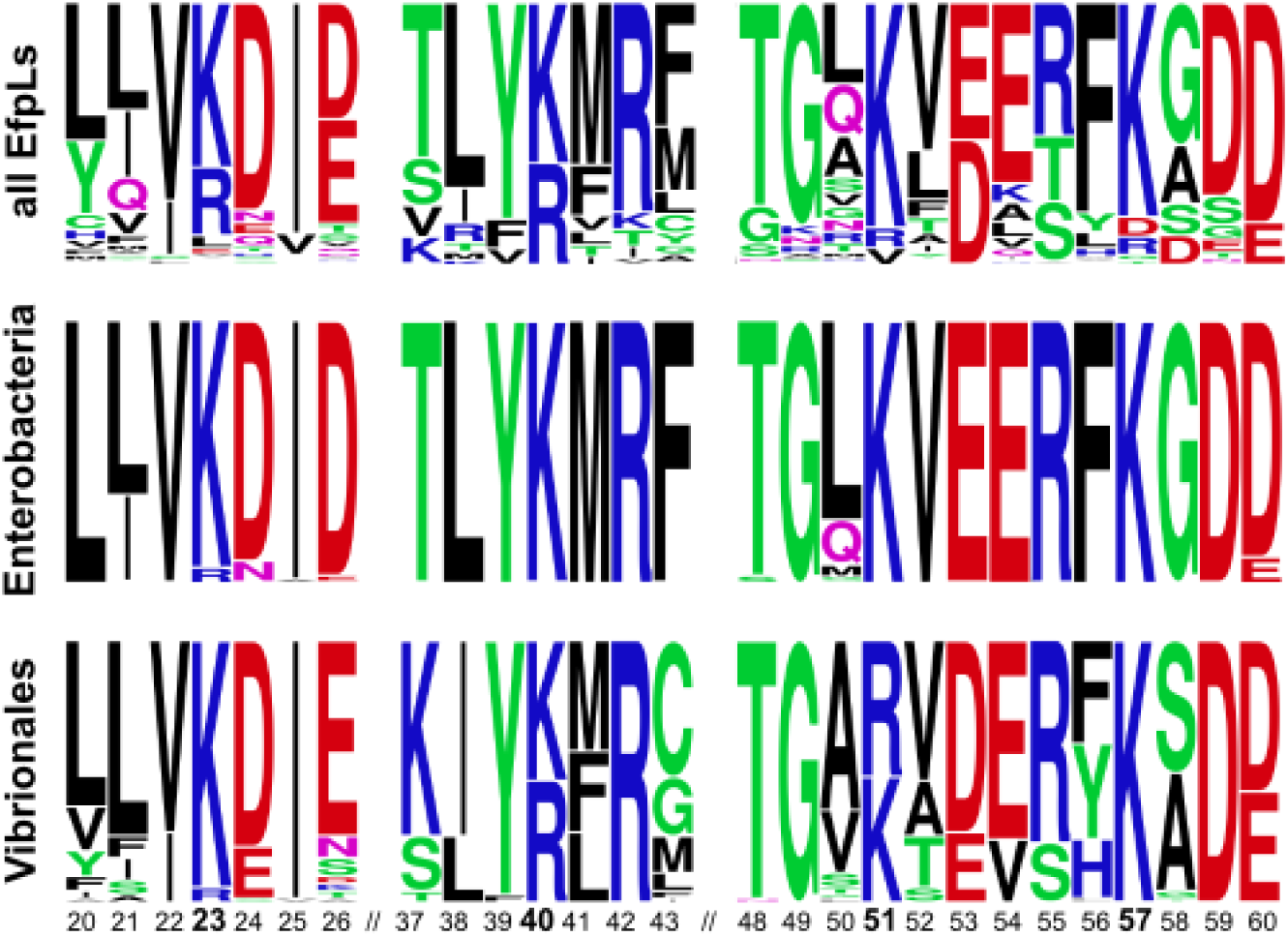
conservation status of acylation sites. Sequence logos^25^ for amino acids at positions 20-60 in all EfpLs, or EfpL from Enterobacteriales or Vibrionales.

**Extended Data Fig. 10:**
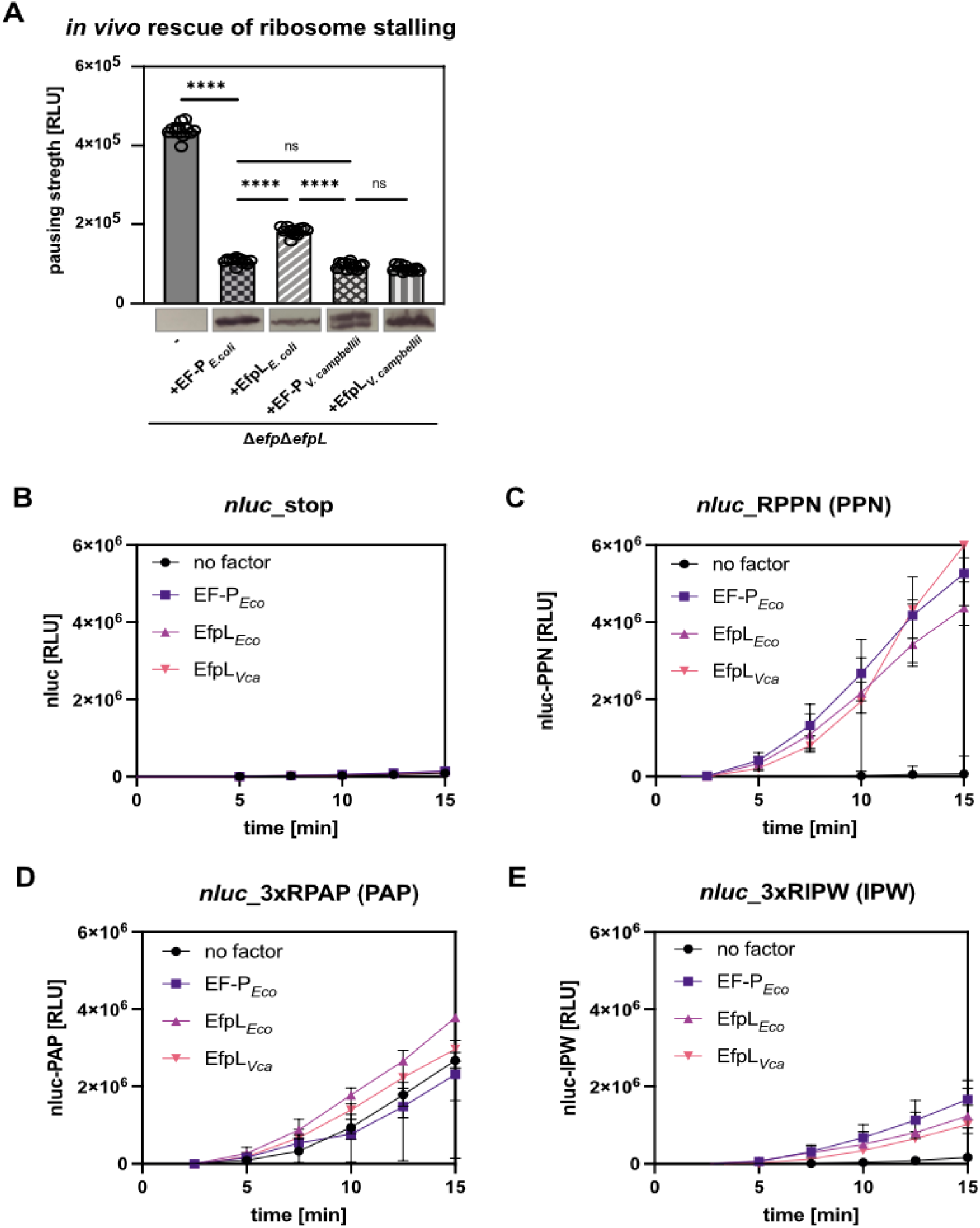
Functional comparison of EfpL from *E. coli* and *V. campbellii*. **(A)** *In vivo* comparison of stalling strength of a PPN motif of *E. coli* Δ*efp*Δ*efpL* cells and respective trans complementation with *E. coli* EF-P (+EF-P*_Eco_*) and EfpL (+EfpL*_Eco_*), as well as *V. campbellii* EF-P (+EF-P*_Vca_*) and EfpL (+EfpL*_Vca_*) (n = 12, Error bars indicate standard deviation). Stalling strength correlates with light emission and is given in relative light units (RLU). Statistically significant differences according to ordinary one-way ANOVA test (*P value <0.0332, **P value <0.0021, ***P value <0.0002, ****P value <0.0001, ns not significant). **(B-E)** *In vitro* transcription and translation of the *nLuc®* variants **(B)** nLuc_stop (no motif), **(C)** nLuc_RPPN (PPN), **(D)** nLuc_3xRIPW (IPW) or **(E)** nLuc_3xRPAP (PAP). The absence (no factor) or presence of the respective translation elongation factors of *E. coli* (EF-P*_Eco_*, EfpL*_Eco_*) or *V. campbellii* (EfpL*_Vca_*) is shown. Translational output was determined by measuring bioluminescence in a time course of 15 minutes (n ≥ 3, Error bars indicate standard deviation).

**Extended Data Fig. 11:**
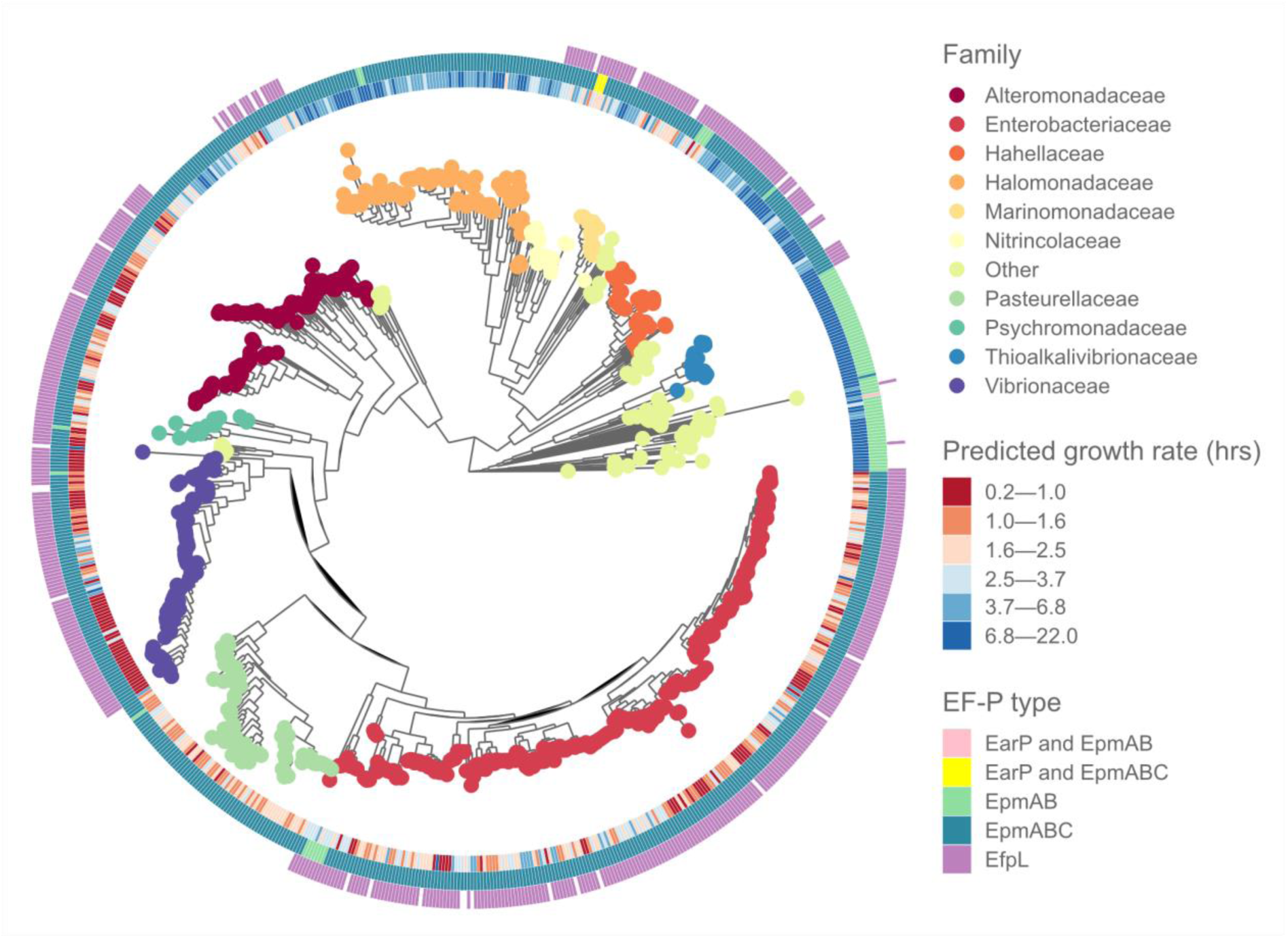
Phylogenetic analysis of predicted growth rates. Set of 920 genomes from the class γ-proteobacteria from the Integrated Microbial Genomes (IMG) database^98^. Inner ring shows doubling times predicted from codon usage bias in ribosomal genes, middle two rings show EF-P types. Colors of tip ends depict phylogenetic family.

## References

1 Tanner, D. R., Cariello, D. A., Woolstenhulme, C. J., Broadbent, M. A. & Buskirk, A. R. Genetic identification of nascent peptides that induce ribosome stalling. J. Biol. Chem. 284, 34809–34818 (2009). 10.1074/jbc.M109.039040

2 Doerfel, L. K. et al. EF-P is essential for rapid synthesis of proteins containing consecutive proline residues. Science 339, 85–88 (2013). 10.1126/science.1229017

3 Ude, S. et al. Translation elongation factor EF-P alleviates ribosome stalling at polyproline stretches. Science 339, 82–85 (2013). 10.1126/science.1228985

4 Hersch, S. J. et al. Divergent protein motifs direct elongation factor P-mediated translational regulation in *Salmonella enterica* and *Escherichia coli*. mBio 4, e00180–00113 (2013). 10.1128/mBio.00180-13

5 Peil, L. et al. Distinct X/PP/X sequence motifs induce ribosome stalling, which is rescued by the translation elongation factor EF-P. Proc. Natl. Acad. Sci. 110, 15265–15270 (2013). 10.1073/pnas.1310642110

6 Woolstenhulme, C. J., Guydosh, N. R., Green, R. & Buskirk, A. R. High-Precision Analysis of Translational Pausing by Ribosome Profiling in Bacteria Lacking EFP. Cell Rep. 11, 13–21 (2015). 10.1016/j.celrep.2015.03.014

7 Gutierrez, E. et al. eIF5A Promotes Translation of Polyproline Motifs. Mol. Cell 51, 1–11 (2013).

8 Lassak, J., Wilson, D. N. & Jung, K. Stall no more at polyproline stretches with the translation elongation factors EF-P and IF-5A. Mol. Microbiol. 99, 219–235 (2016). 10.1111/mmi.13233

9 Katoh, T., Wohlgemuth, I., Nagano, M., Rodnina, M. V. & Suga, H. Essential structural elements in tRNA^Pro^ for EF-P-mediated alleviation of translation stalling. Nat. Commun. 7 (2016). https://doi.org:ARTN 11657 10.1038/ncomms11657

10 Huter, P. et al. Structural Basis for Polyproline-Mediated Ribosome Stalling and Rescue by the Translation Elongation Factor EF-P. Mol. Cell 68, 515–527.e516 (2017). 10.1016/j.molcel.2017.10.014

11 Blaha, G., Stanley, R. E. & Steitz, T. A. Formation of the first peptide bond: the structure of EF-P bound to the 70S ribosome. Science 325, 966–970 (2009). 10.1126/science.1175800

12 Hanawa-Suetsugu, K. et al. Crystal structure of elongation factor P from *Thermus thermophilus* HB8. Proc. Natl. Acad. Sci. 101, 9595–9600 (2004). 10.1073/pnas.03086671010308667101 [pii]

13 Katz, A., Elgamal, S., Rajkovic, A. & Ibba, M. Non-canonical roles of tRNAs and tRNA mimics in bacterial cell biology. Mol. Microbiol. 101, 545–558 (2016). 10.1111/mmi.13419

14 Lassak, J. et al. Arginine-rhamnosylation as new strategy to activate translation elongation factor P. Nat. Chem. Biol. 11, 266–270 (2015). 10.1038/nchembio.1751

15 Bailly, M. & de Crecy-Lagard, V. Predicting the pathway involved in post-translational modification of elongation factor P in a subset of bacterial species. Biol. Direct 5, 3 (2010). https://doi.org:1745-6150-5-3 [pii] 10.1186/1745-6150-5-3

16 Yanagisawa, T., Sumida, T., Ishii, R., Takemoto, C. & Yokoyama, S. A paralog of lysyl-tRNA synthetase aminoacylates a conserved lysine residue in translation elongation factor P. Nat. Struct. Mol. Biol. 17, 1136–1143 (2010). 10.1038/nsmb.1889

17 Navarre, W. W. et al. PoxA, YjeK, and elongation factor P coordinately modulate virulence and drug resistance in *Salmonella enterica*. Mol. Cell 39, 209–221 (2010). https://doi.org:S1097-2765(10)00462-4 [pii] 10.1016/j.molcel.2010.06.021

18 Roy, H. et al. The tRNA synthetase paralog PoxA modifies elongation factor-P with (*R*)-β-lysine. Nat. Chem. Biol. 7, 667–669 (2011). 10.1038/nchembio.632

19 Peil, L. et al. Lys34 of translation elongation factor EF-P is hydroxylated by YfcM. Nat. Chem. Biol. 8, 695–697 (2012). 10.1038/nchembio.1001

20 Rajkovic, A. et al. Translation Control of Swarming Proficiency in *Bacillus subtilis* by 5-Amino-pentanolylated Elongation Factor P. J. Biol. Chem. 291, 10976–10985 (2016). 10.1074/jbc.M115.712091

21 Li, X. et al. Resolving the α-glycosidic linkage of arginine-rhamnosylated translation elongation factor P triggers generation of the first Arg^Rha^ specific antibody. Chem. Sci. 7, 6995–7001 (2016). 10.1039/c6sc02889f

22 Krafczyk, R. et al. Structural Basis for EarP-Mediated Arginine Glycosylation of Translation Elongation Factor EF-P. mBio 8 (2017). 10.1128/mBio.01412-17

23 Brewer, T. E. & Wagner, A. Horizontal gene transfer of a key translation protein has shaped the polyproline proteome. bioRxiv, 2023.2010.2019.563058 (2023). 10.1101/2023.10.19.563058

24 O’Leary, N. A. et al. Reference sequence (RefSeq) database at NCBI: current status, taxonomic expansion, and functional annotation. Nucleic Acids Res. 44, D733–745 (2016). 10.1093/nar/gkv1189

25 Crooks, G. E., Hon, G., Chandonia, J. M. & Brenner, S. E. WebLogo: a sequence logo generator. Genome Res. 14, 1188–1190 (2004). 10.1101/gr.849004

26 Golubev, A. et al. NMR and crystallographic structural studies of the Elongation factor P from *Staphylococcus aureus*. Eur. Biophys. J. 49, 223–230 (2020). 10.1007/s00249-020-01428-x

27 van Zundert, G. C. P. et al. The HADDOCK2.2 Web Server: User-Friendly Integrative Modeling of Biomolecular Complexes. J. Mol. Biol. 428, 720–725 (2016). 10.1016/j.jmb.2015.09.014

28 Schmidt, A. et al. The quantitative and condition-dependent *Escherichia coli* proteome. Nat. Biotechnol. 34, 104–110 (2016). 10.1038/nbt.3418

29 Jung, K., Fabiani, F., Hoyer, E. & Lassak, J. Bacterial transmembrane signalling systems and their engineering for biosensing. Open Biol. 8 (2018). 10.1098/rsob.180023

30 Krafczyk, R. et al. Proline codon pair selection determines ribosome pausing strength and translation efficiency in bacteria. *Commun*. Biol. 4, 589 (2021). 10.1038/s42003-021-02115-z

31 Kumari, R., Michel, A. M. & Baranov, P. V. PausePred and Rfeet: webtools for inferring ribosome pauses and visualizing footprint density from ribosome profiling data. RNA 24, 1297–1304 (2018). 10.1261/rna.065235.117

32 Mi, H. et al. Protocol Update for large-scale genome and gene function analysis with the PANTHER classification system (v.14.0). Nat. Protoc. 14, 703–721 (2019). 10.1038/s41596-019-0128-8

33 Schuller, A. P., Wu, C. C., Dever, T. E., Buskirk, A. R. & Green, R. eIF5A Functions Globally in Translation Elongation and Termination. Mol. Cell 66, 194–205.e195 (2017). 10.1016/j.molcel.2017.03.003

34 Pelechano, V. & Alepuz, P. eIF5A facilitates translation termination globally and promotes the elongation of many non polyproline-specific tripeptide sequences. Nucleic Acids Res. 45, 7326–7338 (2017). 10.1093/nar/gkx479

35 Gall, A. R. et al. Mg^2+^ regulates transcription of *mgtA* in *Salmonella Typhimurium* via translation of proline codons during synthesis of the MgtL peptide. Proc. Natl. Acad. Sci. 113, 15096–15101 (2016). 10.1073/pnas.1612268113

36 Nam, D., Choi, E., Shin, D. & Lee, E. J. tRNA^Pro^-mediated downregulation of elongation factor P is required for *mgtCBR* expression during *Salmonella* infection. Mol. Microbiol. 102, 221–232 (2016). 10.1111/mmi.13454

37 Takada, H., Fujiwara, K., Atkinson, G. C., Chiba, S. & Hauryliuk, V. Resolution of ribosomal stalling by ABCF ATPases YfmR and YkpA/YbiT. bioRxiv, 2024.2001.2025.577322 (2024). 10.1101/2024.01.25.577322

38 Mudryi, V. Elongation factor P: mechanism of action and opportunities for drug design, Georg-August Universität Göttingen, (2023).

39 Qi, F., Motz, M., Jung, K., Lassak, J. & Frishman, D. Evolutionary analysis of polyproline motifs in *Escherichia coli* reveals their regulatory role in translation. PLoS Comput. Biol. 14, e1005987 (2018). 10.1371/journal.pcbi.1005987

40 Mohapatra, S., Choi, H., Ge, X., Sanyal, S. & Weisshaar, J. C. Spatial distribution and ribosome-binding dynamics of EF-P in live *Escherichia coli*. mBio 8 (2017). 10.1128/mBio.00300-17

41 Ishfaq, M. et al. Acetylation regulates subcellular localization of eukaryotic translation initiation factor 5A (eIF5A). FEBS Lett. 586, 3236–3241 (2012). 10.1016/j.febslet.2012.06.042

42 Chung, J., Rocha, A. A., Tonelli, R. R., Castilho, B. A. & Schenkman, S. Eukaryotic initiation factor 5A dephosphorylation is required for translational arrest in stationary phase cells. Biochem. J. 451, 257–267 (2013). 10.1042/bj20121553

43 Kuhn, M. L. et al. Structural, kinetic and proteomic characterization of acetyl phosphate-dependent bacterial protein acetylation. PLoS One 9, e94816 (2014). 10.1371/journal.pone.0094816

44 Weinert, B. T. et al. Acetyl-phosphate is a critical determinant of lysine acetylation in *E. coli*. Mol. Cell 51, 265–272 (2013). 10.1016/j.molcel.2013.06.003

45 Weinert, B. T. et al. Lysine succinylation is a frequently occurring modification in prokaryotes and eukaryotes and extensively overlaps with acetylation. Cell Rep. 4, 842–851 (2013). 10.1016/j.celrep.2013.07.024

46 Qian, L. et al. Global Profiling of Protein Lysine Malonylation in *Escherichia coli* Reveals Its Role in Energy Metabolism. J. Proteome Res. 15, 2060–2071 (2016). 10.1021/acs.jproteome.6b00264

47 Volkwein, W., Maier, C., Krafczyk, R., Jung, K. & Lassak, J. A Versatile Toolbox for the Control of Protein Levels Using *N*^ε^-Acetyl-L-lysine Dependent Amber Suppression. ACS Synth. Biol. 6, 1892–1902 (2017). 10.1021/acssynbio.7b00048

48 Feid, S. C. et al. Regulation of Translation by Lysine Acetylation in *Escherichia coli*. mBio 13, e0122422 (2022). 10.1128/mbio.01224-22

49 Perez, J. C. & Groisman, E. A. Evolution of transcriptional regulatory circuits in bacteria. Cell 138, 233–244 (2009). 10.1016/j.cell.2009.07.002

50 Eagon, R. G. *Pseudomonas natriegens*, a marine bacterium with a generation time of less than 10 minutes. J. Bacteriol. 83, 736–737 (1962). 10.1128/jb.83.4.736-737.1962

51 Weissman, J. L., Hou, S. & Fuhrman, J. A. Estimating maximal microbial growth rates from cultures, metagenomes, and single cells via codon usage patterns. Proc. Natl. Acad. Sci. 118 (2021). 10.1073/pnas.2016810118

52 Starosta, A. L. et al. A conserved proline triplet in Val-tRNA synthetase and the origin of elongation factor P. Cell Rep. 9, 476–483 (2014). 10.1016/j.celrep.2014.09.008

53 Adzhubei, A. A., Sternberg, M. J. & Makarov, A. A. Polyproline-II helix in proteins: structure and function. J. Mol. Biol. 425, 2100–2132 (2013). 10.1016/j.jmb.2013.03.018

54 Pavlov, M. Y. et al. Slow peptide bond formation by proline and other *N*-alkylamino acids in translation. Proc. Natl. Acad. Sci. 106, 50–54 (2009). 10.1073/pnas.0809211106

55 Wohlgemuth, I., Brenner, S., Beringer, M. & Rodnina, M. V. Modulation of the rate of peptidyl transfer on the ribosome by the nature of substrates. J. Biol. Chem. 283, 32229–32235 (2008). 10.1074/jbc.M805316200

56 Mandal, A., Mandal, S. & Park, M. H. Genome-Wide Analyses and Functional Classification of Proline Repeat-Rich Proteins: Potential Role of eIF5A in Eukaryotic Evolution. PLoS One 9, e111800 (2014). 10.1371/journal.pone.0111800

57 Volkwein, W. et al. Switching the post-translational modification of translation elongation factor EF-P. Front. Microbiol. 10, 1148 (2019). 10.3389/fmicb.2019.01148

58 Novozhilov, A. S. & Koonin, E. V. Exceptional error minimization in putative primordial genetic codes. Biol. Direct 4, 44 (2009). 10.1186/1745-6150-4-44

59 Chadani, Y., Uemura, E., Yamazaki, K., Kurihara, M. & Taguchi, H. The ABCF proteins in *Escherichia coli* individually alleviate nascent peptide-induced noncanonical translations. bioRxiv, 2023.2010.2004.560807 (2023). 10.1101/2023.10.04.560807

60 Hong, H.-R., Prince, C. R., Tetreault, D. D., Wu, L. & Feaga, H. A. YfmR is a translation factor that prevents ribosome stalling and cell death in the absence of EF-P. bioRxiv, 2023.2008.2004.552005 (2023). 10.1101/2023.08.04.552005

61 Singh, S. et al. Cryo-EM studies of the four *E. coli* paralogs establish ABCF proteins as master plumbers of the peptidyl-transferase center of the ribosome. bioRxiv, 2023.2006.2015.543498 (2023). 10.1101/2023.06.15.543498

62 Pinheiro, D. F. B. The translation elongation factor P in actinobacteria, Ludwig-Maximilians-Universität München, (2020).

63 Lassak, J., Sieber, A. & Hellwig, M. Exceptionally versatile take II: post-translational modifications of lysine and their impact on bacterial physiology. Biol. Chem. (2022). 10.1515/hsz-2021-0382

64 Veening, J. W. et al. Bet-hedging and epigenetic inheritance in bacterial cell development. Proc. Natl. Acad. Sci. 105, 4393–4398 (2008). 10.1073/pnas.0700463105

65 Hummels, K. R. & Kearns, D. B. Translation elongation factor P (EF-P). FEMS Microbiol. Rev. 44, 208–218 (2020). 10.1093/femsre/fuaa003

66 Pinheiro, B. et al. Structure and Function of an Elongation Factor P Subfamily in Actinobacteria. Cell Rep. 30, 4332–4342.e4335 (2020). 10.1016/j.celrep.2020.03.009

67 Doerfel, L. K. et al. Entropic Contribution of Elongation Factor P to Proline Positioning at the Catalytic Center of the Ribosome. J. Am. Chem. Soc. 137, 12997–13006 (2015). 10.1021/jacs.5b07427

68 Katoh, T., Tajima, K. & Suga, H. Consecutive elongation of D-amino acids in translation. Cell Chem. Biol. 24, 46–54 (2017). 10.1016/j.chembiol.2016.11.012

69 Katoh, T. & Suga, H. Ribosomal incorporation of consecutive β-amino acids. J. Am. Chem. Soc. 140, 12159–12167 (2018). 10.1021/jacs.8b07247

70 Katoh, T. & Suga, H. Translation initiation with exotic amino acids using EF-P-responsive artificial initiator tRNA. Nucleic Acids Res. 51, 8169–8180 (2023). 10.1093/nar/gkad496

71 Guzman, L. M., Belin, D., Carson, M. J. & Beckwith, J. Tight regulation, modulation, and high-level expression by vectors containing the arabinose P_BAD_ promoter. J. Bacteriol. 177, 4121–4130 (1995). 10.1128/jb.177.14.4121-4130.1995

72 Lassak, J., Henche, A. L., Binnenkade, L. & Thormann, K. M. ArcS, the cognate sensor kinase in an atypical Arc system of *Shewanella oneidensis* MR-1. Appl. Environ. Microbiol. 76, 3263–3274 (2010). 10.1128/aem.00512-10

73 Lassak, J., Bubendorfer, S. & Thormann, K. M. Domain analysis of ArcS, the hybrid sensor kinase of the *Shewanella oneidensis* MR-1 Arc two-component system, reveals functional differentiation of its two receiver domains. J. Bacteriol. 195, 482–492 (2013). 10.1128/jb.01715-12

74 Gödeke, J., Heun, M., Bubendorfer, S., Paul, K. & Thormann, K. M. Roles of two *Shewanella oneidensis* MR-1 extracellular endonucleases. Appl. Environ. Microbiol. 77, 5342–5351 (2011). 10.1128/aem.00643-11

75 Bertani, G. Studies on lysogenesis. I. The mode of phage liberation by lysogenic *Escherichia coli*. J. Bacteriol. 62, 293–300 (1951). 10.1128/jb.62.3.293-300.1951

76 Miller, J. H. Experiments in molecular genetics. (Cold Spring Harbor Laboratory 1972).

77 Hanahan, D. Studies on transformation of *Escherichia coli* with plasmids. J. Mol. Biol. 166, 557–580 (1983). 10.1016/s0022-2836(83)80284-8

78 Nandi, B. et al. Rapid method for species-specific identification of *Vibrio cholerae* using primers targeted to the gene of outer membrane protein OmpW. J. Clin. Microbiol. 38, 4145–4151 (2000). 10.1128/jcm.38.11.4145-4151.2000

79 Gallego-Jara, J., Écija Conesa, A., de Diego Puente, T., Lozano Terol, G. & Cánovas Díaz, M. Characterization of CobB kinetics and inhibition by nicotinamide. PLoS One 12, e0189689 (2017). 10.1371/journal.pone.0189689

80 Laemmli, U. K. Cleavage of structural proteins during the assembly of the head of bacteriophage T4. Nature 227, 680–685 (1970). 10.1038/227680a0

81 Ladner, C. L., Yang, J., Turner, R. J. & Edwards, R. A. Visible fluorescent detection of proteins in polyacrylamide gels without staining. Anal. Biochem. 326, 13–20 (2004). 10.1016/j.ab.2003.10.047

82 Martin, M. Cutadapt removes adapter sequences from high-throughput sequencing reads. EMBnet 17, 3 (2011). 10.14806/ej.17.1.200

83 Langmead, B., Trapnell, C., Pop, M. & Salzberg, S. L. Ultrafast and memory-efficient alignment of short DNA sequences to the human genome. Genome Biol. 10, R25 (2009). 10.1186/gb-2009-10-3-r25

84 Finn, R. D. et al. The Pfam protein families database: towards a more sustainable future. Nucleic Acids Res. 44, D279–285 (2016). 10.1093/nar/gkv1344

85 Sievers, F. & Higgins, D. G. The Clustal Omega Multiple Alignment Package. Methods Mol. Biol. 2231, 3–16 (2021). 10.1007/978-1-0716-1036-7_1

86 Price, M. N., Dehal, P. S. & Arkin, A. P. FastTree 2--approximately maximum-likelihood trees for large alignments. PLoS One 5, e9490 (2010). 10.1371/journal.pone.0009490

87 Yu, G., Smith, D. K., Zhu, H., Guan, Y. & Lam, T. T.-Y. ggtree: an r package for visualization and annotation of phylogenetic trees with their covariates and other associated data. Methods Ecol. Evol. 8, 28–36 (2017). 10.1111/2041-210X.12628

88 Cianci, M. et al. P13, the EMBL macromolecular crystallography beamline at the low-emittance PETRA III ring for high- and low-energy phasing with variable beam focusing. J. Synchrotron. Radiat. 24, 323–332 (2017). 10.1107/s1600577516016465

89 Vonrhein, C. et al. Advances in automated data analysis and processing within autoPROC, combined with improved characterisation, mitigation and visualisation of the anisotropy of diffraction limits using STARANISO. Acta Crystallogr. A Found. Adv. (2018).

90 Krissinel, E., Uski, V., Lebedev, A., Winn, M. & Ballard, C. Distributed computing for macromolecular crystallography. Acta Crystallogr. D Struct. Biol. 74, 143–151 (2018). 10.1107/s2059798317014565

91 Jumper, J. et al. Highly accurate protein structure prediction with AlphaFold. Nature 596, 583–589 (2021). 10.1038/s41586-021-03819-2

92 Varadi, M. et al. AlphaFold Protein Structure Database: massively expanding the structural coverage of protein-sequence space with high-accuracy models. Nucleic Acids Res. 50, D439–d444 (2022). 10.1093/nar/gkab1061

93 Bond, P. S. & Cowtan, K. D. ModelCraft: an advanced automated model-building pipeline using Buccaneer. Acta Crystallogr. D Struct. Biol. 78, 1090–1098 (2022). 10.1107/s2059798322007732

94 Joosten, R. P., Long, F., Murshudov, G. N. & Perrakis, A. The PDB_REDO server for macromolecular structure model optimization. IUCrJ 1, 213–220 (2014). 10.1107/s2052252514009324

95 Kovalevskiy, O., Nicholls, R. A., Long, F., Carlon, A. & Murshudov, G. N. Overview of refinement procedures within REFMAC5: utilizing data from different sources. Acta Crystallogr. D Struct. Biol. 74, 215–227 (2018). 10.1107/s2059798318000979

96 Emsley, P., Lohkamp, B., Scott, W. G. & Cowtan, K. Features and development of Coot. Acta Crystallogr. D Biol. Crystallogr. 66, 486–501 (2010). 10.1107/s0907444910007493

97 Williams, C. J. et al. MolProbity: More and better reference data for improved all-atom structure validation. Protein Sci. 27, 293–315 (2018). 10.1002/pro.3330

98 Chen, I. A. et al. The IMG/M data management and analysis system v.6.0: new tools and advanced capabilities. Nucleic Acids Res. 49, D751–d763 (2021). 10.1093/nar/gkaa939

99 Parks, D. H., Imelfort, M., Skennerton, C. T., Hugenholtz, P. & Tyson, G. W. CheckM: assessing the quality of microbial genomes recovered from isolates, single cells, and metagenomes. Genome Res. 25, 1043–1055 (2015). 10.1101/gr.186072.114

100 Chaumeil, P.-A., Mussig, A. J., Hugenholtz, P. & Parks, D. H. GTDB-Tk: a toolkit to classify genomes with the Genome Taxonomy Database. Bioinformatics 36, 1925–1927 (2019). 10.1093/bioinformatics/btz848

101 Galperin, M. Y. et al. COG database update: focus on microbial diversity, model organisms, and widespread pathogens. Nucleic Acids Res. 49, D274–D281 (2020). 10.1093/nar/gkaa1018

102 Mistry, J. et al. Pfam: The protein families database in 2021. Nucleic Acids Res. 49, D412–d419 (2021). 10.1093/nar/gkaa913

103 Li, W. et al. RefSeq: expanding the Prokaryotic Genome Annotation Pipeline reach with protein family model curation. Nucleic Acids Res. 49, D1020–d1028 (2021). 10.1093/nar/gkaa1105

104 Revell, L. J. phytools: an R package for phylogenetic comparative biology (and other things). Methods Ecol. Evol. 3, 217–223 (2012). 10.1111/j.2041-210X.2011.00169.x

105 Edgar, R. C. MUSCLE: multiple sequence alignment with high accuracy and high throughput. Nucleic Acids Res. 32, 1792–1797 (2004). 10.1093/nar/gkh340

106 Nguyen, L. T., Schmidt, H. A., von Haeseler, A. & Minh, B. Q. IQ-TREE: a fast and effective stochastic algorithm for estimating maximum-likelihood phylogenies. Mol. Biol. Evol. 32, 268–274 (2015). 10.1093/molbev/msu300

107 Kalyaanamoorthy, S., Minh, B. Q., Wong, T. K. F., von Haeseler, A. & Jermiin, L. S. ModelFinder: fast model selection for accurate phylogenetic estimates. Nat. Methods 14, 587–589 (2017). 10.1038/nmeth.4285

108 Wickham, H. Ggplot2 : elegant graphics for data analysis. (Springer Science+Business Media, LLC, 2016).

