## Supplementary Figures for "EF-P and its paralog EfpL (YeiP) differentially control translation of proline-containing sequences"

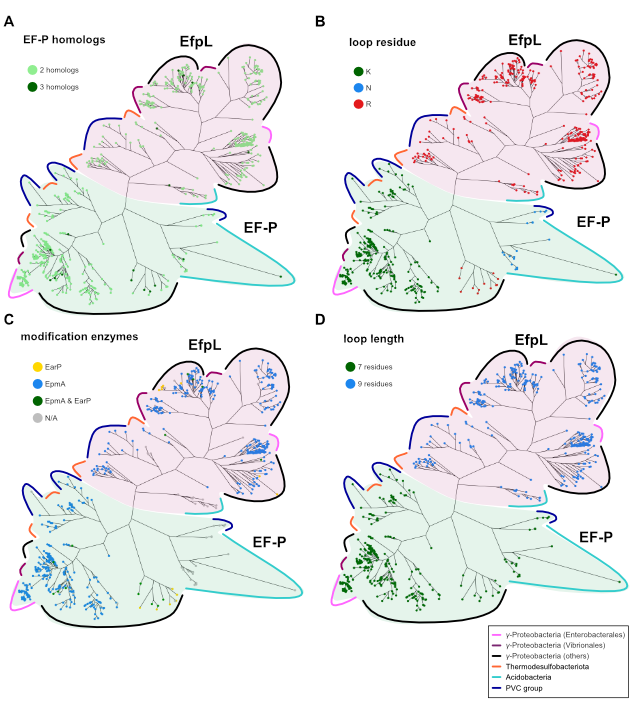

**Supplementary Fig. S1: Phylogenetic analysis of co-occurring EF-P and EfpL proteins**

Phylogenetic tree of EfpL (purple) and co-occurring EF-Ps (green). Colored lines indicate bacterial phyla. **(A)** EF-P homologs per proteome. **(B)** β3Ωβ4 loop tip residue in EF-P or EfpL. **(C)** EF-P modification enzymes found per proteome. **(D)** β3Ωβ4 loop length of EF-P or EfpL.

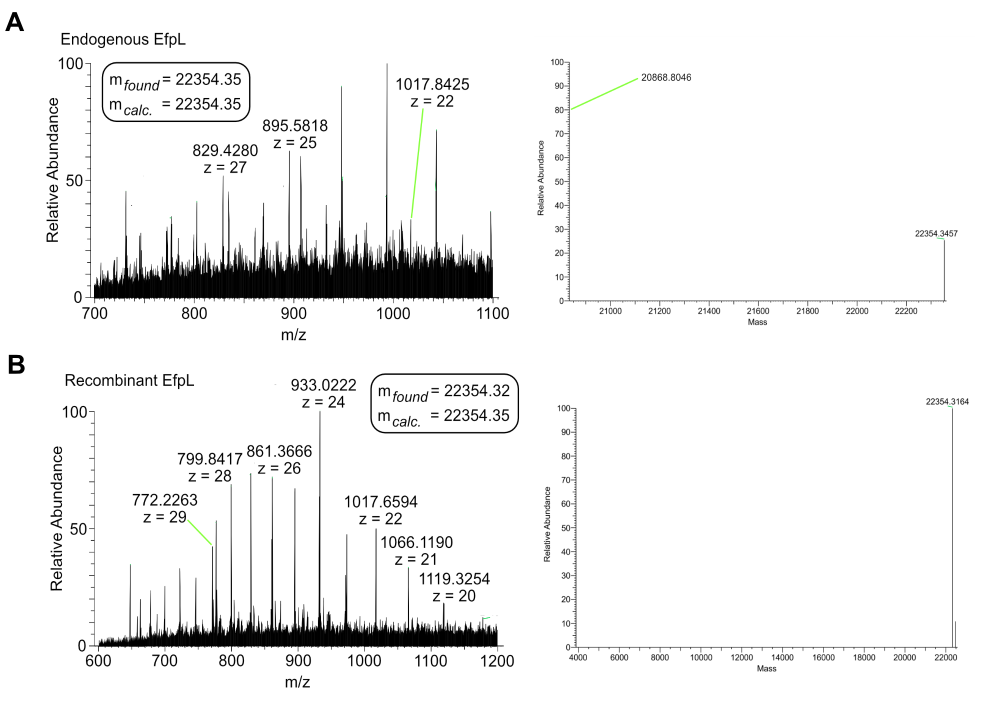

**Supplementary Fig. S2: Mass spectrometry (MS) for EfpL protein analysis**

MS spectra for **(A)** endogeneous *E. coli* EfpL and **(B)** recombinant produced *E. coli* EfpL to identify modification status. Left side: DDA raw files; right side: output mzML format. Mass (m*_calc._*) was calculated according the Uniprot database (identifier: B7UFI8 - EFPL_ECO27)^1^. At least two unique peptides were required for protein identification. False discovery rate determination was carried out using a decoy database and thresholds were set to 1 % FDR.

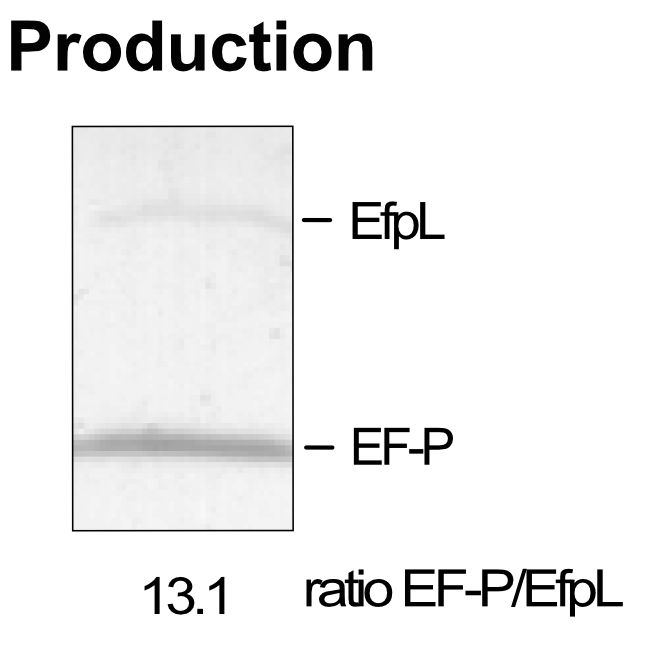

**Supplementary Fig. S3: Protein amount of EF-P and EfpL in *E. coli***

To quantify endogenous production of EF-P and EfpL a 6xHis encoding sequence was genomically integrated at the 3' end of the ORFs of *efp* and *efpL* in *E. coli* BW25113. Production was quantified via immunoblotting using Anti-His6 antibodies. Ratio determined using Fiji^2^.

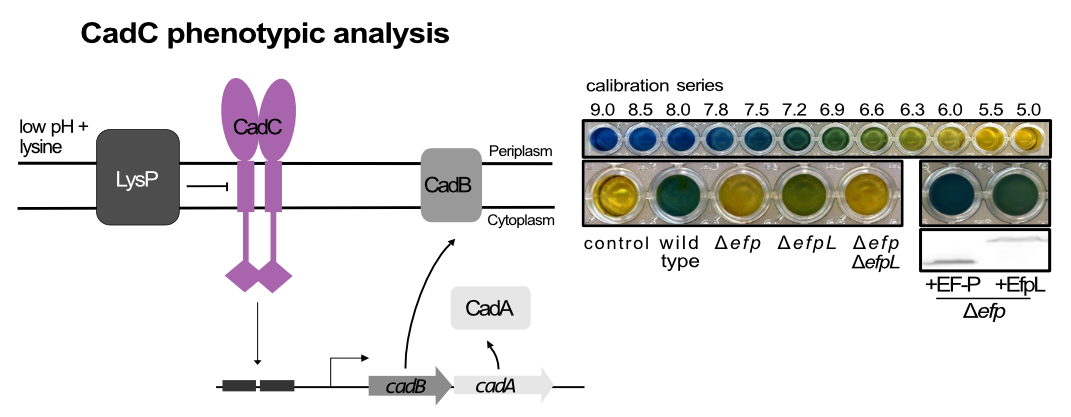

**Supplementary Fig. S4: CadC phenotypic analysis**

Scheme of CadC dependent pH regulation^3^: To visualize pH regulation, cells were cultivated in lysine deccarboxylase indicator medium (indicator: bromothymol blue) and alkalization is depicted as a color change from yellow over green to blue. Production of EF-P and EfpL was confirmed by immunodetection of the C-terminally attached His6-tag using Anti-His6 antibodies.

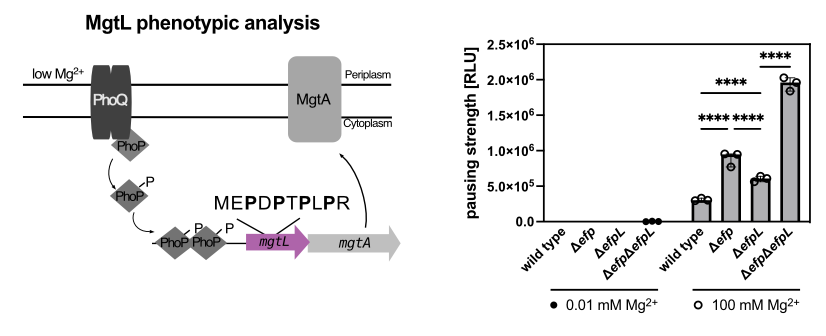

**Supplementary Fig. S5: MgtL phenotypic analysis**

Left: model illustrating the regulation mechanism of Mg^2+^ uptake by MgtA ^4-6^. *mgtL* consists of a proline-rich sequence and regulates *mgtA* expression. Right: reporter assay to detect pausing strength at the MgtL leader peptide with the sequence MEPDPTPLPR. Maximal luminescence emission under high (100 mM) and low (100 µM) Mg^2+^ in *E. coli* BW23113 and corresponding mutant strains is depicted (n = 12, Error bars indicate standard deviation). Pausing strength correlates with light emission and is given in relative light units (RLU). Statistically significant differences according to two-sided t-test (*P value <0.0332, **P value <0.0021, ***P value <0.0002, ****P value <0.0001, ns not significant).

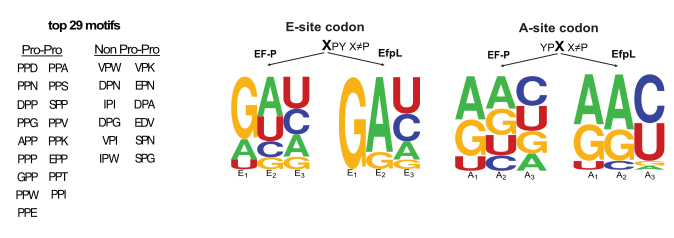

**Supplementary Fig. S6: Top 29 motifs in *E. coli* and comparison of codon bias of the E-site and A-site amino acid**

**(A)** Top 29 EF-P dependent arrest motifs associated with ribosome pausing in *E. coli* BW25113 determined by PausePred^7^. **(B)** Sequence logos of the E- and A-site codons in XPY or YPX arrest motifs X≠P (24) targeted by EF-P and EfpL, respectively.

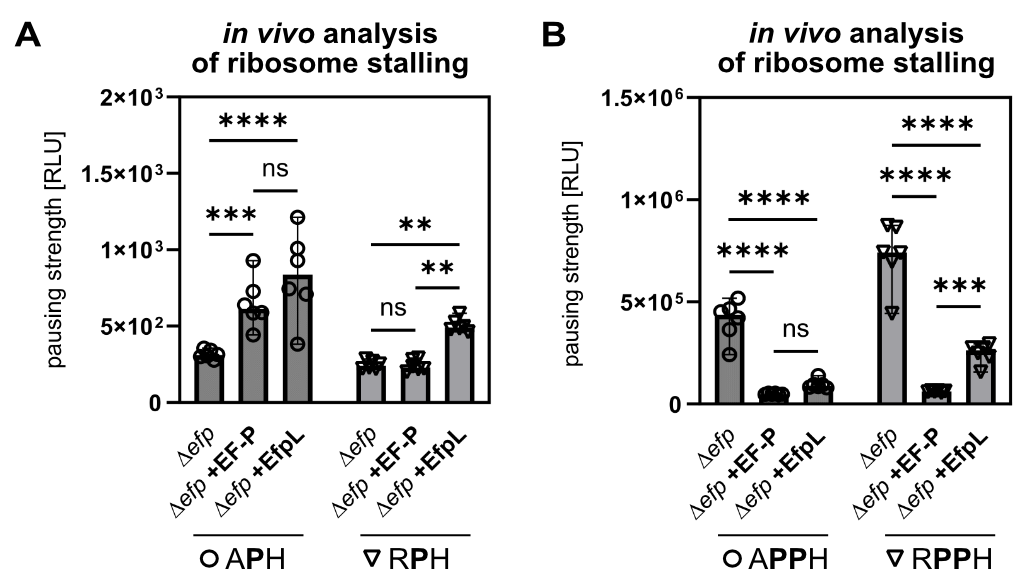

**Supplementary Fig. S7: EF-P and EfpL can induce ribosome stalling**

*In vivo* comparison of stalling strength of a **(A)** APH and RPH or **(B)** APPH and RPPH motif in Δ*efp* strains in the absence or presence of *efp* (+EF-P), or *efpL* (+EfpL). Pausing strength correlates with light emission and is given in relative light units (RLU) (n = 12, Error bars indicate standard deviation). Statistically significant differences according to ordinary one-way ANOVA test (*P value <0.0332, **P value <0.0021, ***P value <0.0002, ****P value <0.0001, ns not significant).

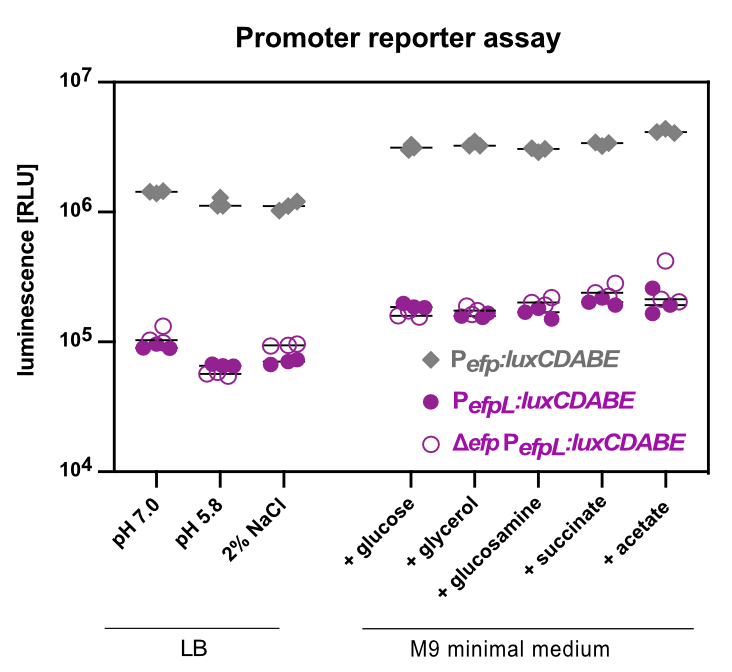

**Supplementary Fig. S8: *efp* and *efpL* gene expression analysis under certain conditions**

Promoters of *efp* (P*_efp_*) or *efpL* (P*_efpL_*) were fused with the *luxCDABE* genes of *P. luminescens* and tested under different conditions. *E. coli* wild type or Δ*efp* strains were transformed with promoter reporter fusions. Maximal luminescence of a 16h time course in LB or M9 minimal medium is given in relative light units (RLU) (n=3, line identifies mean value).

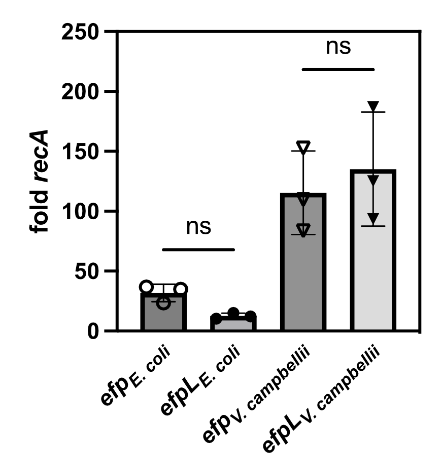

**Supplementary Fig. S9: Expression analysis of EF-P and *EfpL* from *E. coli* and *V. campbellii***

Quantitative real time PCR (qRT-PCRs) were performed to analyze expression of *efp* and *efpL* in *E. coli* or *V. campbellii.* (n=3, Error bars indicate standard deviation). Statistically significant differences according to ordinary one-way ANOVA test (*P value <0.0332, **P value <0.0021, ***P value <0.0002, ****P value <0.0001, ns not significant). Primer efficiency were as following: *recA_E. coli_* 1.987, *efp_E. Coli_* 1.953, *efpL_E. coli_* 1.936, *recA_V. campbellii_* 2.084, *efp_V. campbellii_* 1.962, *efpL_V. campbellii_* 2.009. Normalization with reference gene *recA* for comparison of *efp* and *efpL* expression.

**References**

1	Consortium, T. U. UniProt: the Universal Protein Knowledgebase in 2023. *Nucleic Acids Res.* **51**, D523-D531 (2022). <https://doi.org:10.1093/nar/gkac1052>

2	Schindelin, J. *et al.* Fiji: an open-source platform for biological-image analysis. *Nat. Methods* **9**, 676-682 (2012). <https://doi.org:10.1038/nmeth.2019>

3	Ude, S. *et al.* Translation elongation factor EF-P alleviates ribosome stalling at polyproline stretches. *Science* **339**, 82-85 (2013). <https://doi.org:10.1126/science.1228985>

4	Gall, A. R. *et al.* Mg2+ regulates transcription of mgtA in *Salmonella Typhimurium* via translation of proline codons during synthesis of the MgtL peptide. *Proceedings of the National Academy of Sciences of the United States of America* **113**, 15096-15101 (2016). <https://doi.org:10.1073/pnas.1612268113>

5	Nam, D., Choi, E., Shin, D. & Lee, E. J. tRNA(Pro) -mediated downregulation of elongation factor P is required for mgtCBR expression during *Salmonella* infection. *Mol. Microbiol.* **102**, 221-232 (2016). <https://doi.org:10.1111/mmi.13454>

6	Takada, H., Fujiwara, K., Atkinson, G. C., Chiba, S. & Hauryliuk, V. Resolution of ribosomal stalling by ABCF ATPases YfmR and YkpA/YbiT. *bioRxiv*, 2024.2001.2025.577322 (2024). <https://doi.org:10.1101/2024.01.25.577322>

7	Kumari, R., Michel, A. M. & Baranov, P. V. PausePred and Rfeet: webtools for inferring ribosome pauses and visualizing footprint density from ribosome profiling data. *RNA* **24**, 1297-1304 (2018). <https://doi.org:10.1261/rna.065235.117>
